# Microglia-derived Small Extracellular Vesicles Promote Bone Marrow Monocyte Expansion and Recruitment in Glioblastoma

**DOI:** 10.64898/2026.10.12.757688

**Authors:** Isabela Albuja, Miguel Quiralte, Neeharika Panda, Abigail Macaraeg, Minjeong Yeon, Qin Liu, Andrew V. Kossenkov, Thibaut Barnoud, Irene Bertolini

## Abstract

Glioblastoma (GBM) remains one of the deadliest human cancers and is characterized by a profoundly immunosuppressive tumor microenvironment (TME). Although tumor-associated microglia (TAMg) promote GBM progression, their role in regulating immune cell recruitment remains poorly understood. Here, we demonstrate that TAMg-derived small extracellular vesicles (sEVs) mediate bone marrow-derived monocyte recruitment into the GBM TME. We find that these sEVs traffic to the bone marrow, where they selectively target monocytes and their progenitors. Mechanistically, sEV internalization activates MAPK signaling and inflammatory transcriptional programs, driving the differentiation and expansion of inflammatory monocytes. Together, our findings establish microglia-derived sEVs as a previously unrecognized mechanism of communication between the GBM TME and the peripheral immune system, thereby promoting the establishment of an immunosuppressive TME that drives GBM progression.

## INTRODUCTION

Glioblastoma (GBM), the most common and aggressive primary brain tumor, remains one of the deadliest human cancers despite decades of intensive research and therapeutic development^1,2^. The remarkable molecular and cellular complexity of GBM continues to impede the development of effective therapies. Consequently, the standard of care for GBM, surgical resection followed by radiotherapy and chemotherapy, has remained largely unchanged, with modest improvements in patient survival^3–5^. GBM is characterized by a complex, highly immunosuppressive tumor microenvironment (TME)^6^, with extensive infiltration of bone marrow-derived myeloid cells^7^. Bone marrow-derived macrophages and resident microglia, constitute up to 40% of the total tumor mass, and their acquisition of an immunosuppressive phenotype is associated with poor clinical outcomes in GBM^8–10^.

Infiltrating macrophages in the GBM TME originate from bone marrow-derived monocytes, while brain-resident microglia stem from the yolk sac^7^. During tumorigenesis, resident microglia acquire a pro-tumorigenic phenotype and become tumor-associated microglia (TAMg) that promote tumor growth, immune escape, and angiogenesis^11^, in part through the release of cytokines, chemokines, and growth factors^12,13^. During the early stages of gliomagenesis, the immune population in the brain is primarily composed of resident microglia, whereas in later stages, the immune compartment is increasingly dominated by bone marrow-derived myeloid cells^8,14^. Inflammatory monocytes are mobilized from the bone marrow into the circulation. Once in circulation, these monocytes can infiltrate the tumor and differentiate into tumor-associated macrophages. Once within the TME, infiltrating macrophages acquire an immunosuppressive phenotype that supports tumor progression^15^. To date, the role of microglia in the recruitment of bone marrow-derived monocytes remains unclear.

Extracellular vesicles (EVs) are key players in intercellular communication in the tumor microenvironment. EVs are lipid-bound secretory vesicles released by all types of cells, both in physiological and pathological conditions^16,17^. Small extracellular vesicles (sEVs) are defined as EVs with a diameter between 50 and 200 nm. sEVs include both exosomes (endosomal origin) and ectosomes (plasma membrane origin)^18,19^. In GBM, the transfer of sEVs between tumor cells and stromal cells has been shown to regulate immune cell behavior and angiogenesis^20–22^. Here, we sought to determine whether microglia-derived sEVs mediate communication between the tumor and the bone marrow to regulate monocyte recruitment and promote an immunosuppressive TME in GBM.

## RESULTS

### Microglia-derived small extracellular vesicles promote tumor growth and reshape the GBM immune microenvironment

The role of microglia-derived small extracellular vesicles (sEVs) in GBM is still poorly understood. To investigate whether tumor-associated microglia (TAMg)-derived sEVs promote GBM progression, we purified sEVs from activated or resting microglia and evaluated their effects on tumor growth and immune infiltration using two complementary murine glioma models, CT2a and GL261 (**Figure 1A**). CT2a exhibits a mesenchymal-like, PTEN-deficient phenotype, whereas GL261 is more immune-active and less mesenchymal^27,28^. Given the molecular and phenotypic heterogeneity of GBM, these complementary models provide a robust platform to assess the generalizability of our findings. To generate TAMg-derived sEVs, we used two complementary sources of activated microglia: BV2 cells stimulated with glioma-conditioned medium and primary microglia isolated from tumor-bearing mice (**Supplementary Figure 1A-B**). Resting BV2 cells and primary microglia isolated from naïve mice served as controls. Purified microglia exhibited >85% purity and expressed activation markers following tumor stimulation (**Supplementary Figure 1C-F and Supplementary Table 1**). sEV preparations met MISEV criteria, displaying comparable size distributions, concentrations, zeta potentials, tetraspanin expression, and morphology, while lacking contaminating cellular proteins (**Supplementary Figure 2A-J and Supplementary Original Western Blot 1-3**)^18^.

**Figure 1:**
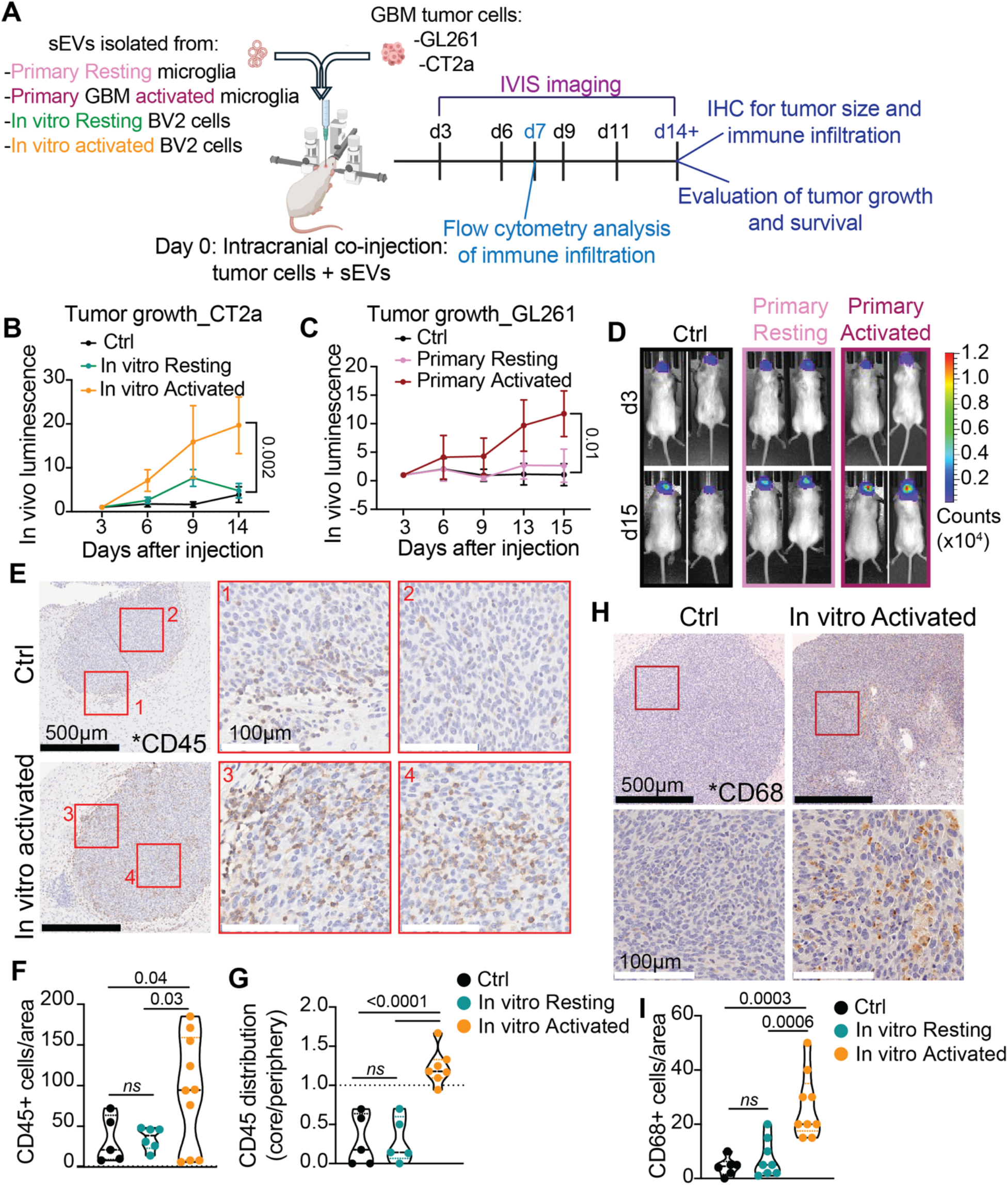
Tumor associated microglia (TAMg)-sEVs accelerate tumor growth. **(A)** Schematic representation of in vivo experiment and experimental readouts. **(B)** CT2a glioma cells were co-injected with BV2-derived sEVs in C57BL/6 male and female albino mice. Tumor growth was tracked using an IVIS SpectrumCT In Vivo Imaging System at 3, 6, 9, and 14 days after injection. Ctrl = mice injected with CT2a cells. In vitro Resting = mice co-injected with CT2a and resting BV2-derived sEVs. In vitro Activated = mice co-injected with CT2a and sEVs derived from BV2 cells activated with CT2a tumor supernatant. n= 8. SD. Two-way ANOVA for repeated measurements. Significance at day 14: p-value = Activated vs Ctrl. Ctrl vs Resting = not significant. Activated vs Resting = p-value 0.0041. Significance at day 9: Activated vs Ctrl = p-value 0.006. Activated vs Resting and Ctrl vs Resting = not significant. **(C)** GL261 glioma cells were co-injected with primary microglia-derived sEVs in C57BL/6 female albino mice. Tumor growth was tracked as in (A). Ctrl = mice injected with GL261 cells. Primary Resting = mice co-injected with GL261 and sEVs isolated from primary microglia isolated from naïve mice. Primary Activated = mice co-injected with GL261 and sEVs produced by primary microglia isolated from mice injected with GL261 cells. n= 6. SD. p-value = Two-way ANOVA for repeated measurements. Significance at day 15. p-value = Ctrl vs Activated. Ctrl vs Resting = not significant. Activated vs Resting = p-value 0.02. **(D)** The conditions are in (C). Representative IVIS imaging of co-injected mice at day 3 (top) and day 15 (bottom). (**E-G**) CT2a glioma cells were co-injected with BV2-derived sEVs in C57BL/6 male and female albino mice. (E) Representative immunohistochemistry images of tumor collected 14 days after the injection and stained for CD45 (immune cells). 4X objective, scale bar = 500μm. Red box = inset, 20X objective, scale bar = 100μm. (F) Quantification of CD45+ cells/area and (G) enrichment of CD45+ cells in the core of the tumor compared to the periphery. One-way ANOVA with Dunnett’s multiple comparisons test. n = 5. (**H-I**) The conditions are as in (B), and tumors were stained for CD68 (macrophages). (H) Representative images. 4X objective, scale bar = 500μm. Red box = inset, 20X objective, scale bar = 100μm. (I) Quantification of CD68+ cells per area. One-way ANOVA with Tukey’s multiple comparisons test. n = 5.

To assess whether microglia-derived sEVs impact tumor growth and the immune microenvironment, luciferase-expressing CT2a or GL261 cells were co-injected intracranially with sEVs isolated from resting or activated microglia (**Figure 1A**). Remarkably, TAMg-derived sEVs significantly increased tumor growth in both glioma models, as assessed by IVIS imaging and histological analysis (**Figure 1B-D; Supplementary Figure 3A-F**).

### scRNAseq analysis reveals that activated microglia-derived sEVs promote monocyte recruitment and immunosuppressive macrophage differentiation

To evaluate the contribution of activated microglia-sEVs to changes the tumor immune infiltrate, first we tested for changes in the infiltration of total CD45+ cells and CD68+ macrophages by immunohistochemistry. Our data showed that the co-injection of TAMg-sEVs with CT2a tumor cells significantly increased the number of infiltrating CD45-positive immune cells (Figure 1E-G and Suppl. Figure 3G) and CD68-positive macrophages (Figure 1H-I and Suppl. Figure 3H).

To define how TAMg-derived sEVs reshape the GBM TME, we performed single-cell RNA sequencing of CD45+ immune cells isolated from GL261 tumors seven days after co-injection with primary microglia-derived sEVs (**Figure 2A**). At this early time point, tumor size remained comparable across experimental groups, allowing immune changes to be evaluated independently of differences in tumor burden. Following quality control filtering (>10,000 cells per sample; **Supplementary Figure 4A**), integrated analysis identified 18 clusters representing eight major immune cell populations (**Figure 2B** and **Supplementary Figure 4B; Supplementary Table 2**).

**Figure 2.**
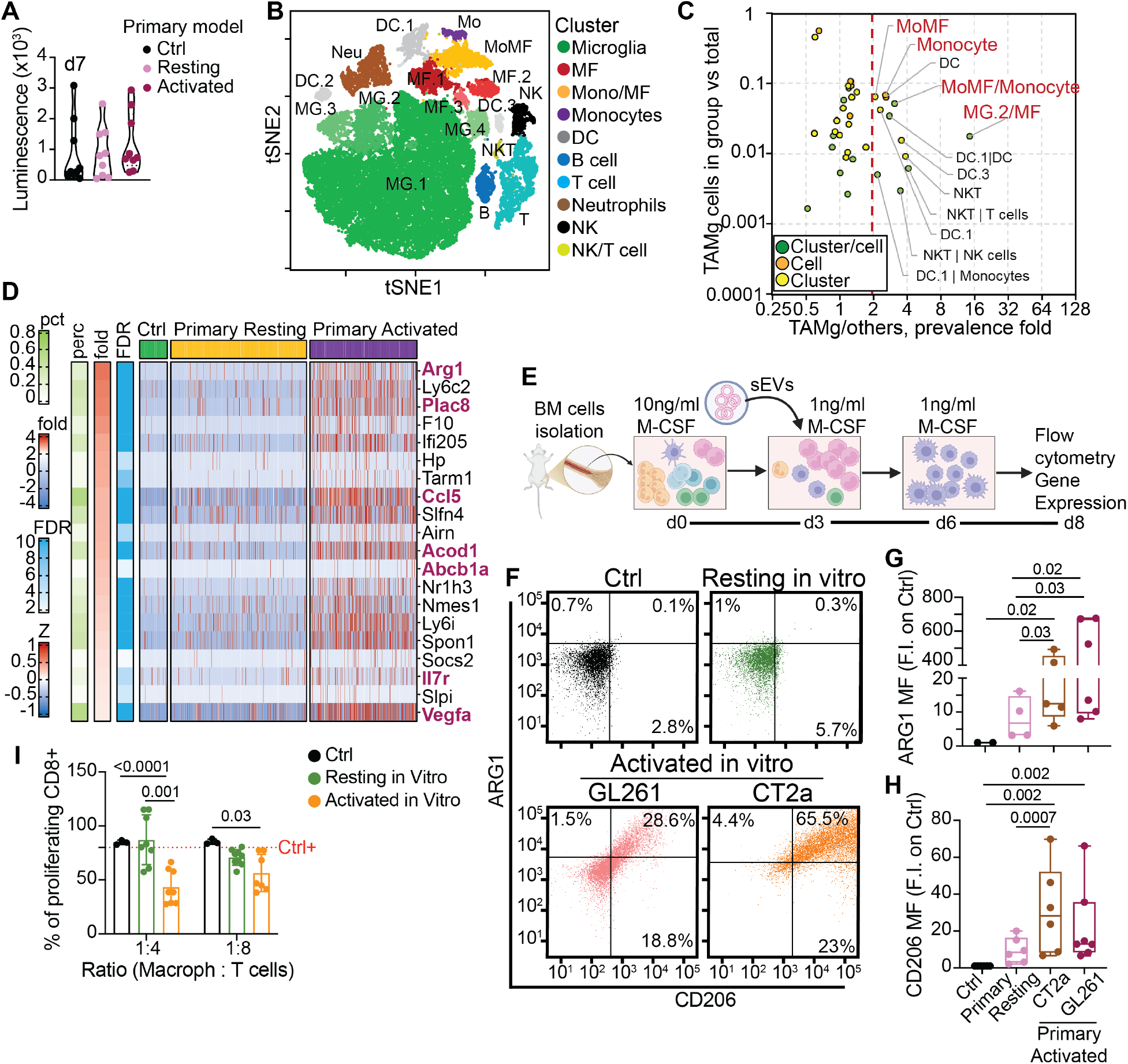
Single cell RNAseq analysis revealed an increase in the infiltration of monocytes and macrophages to the tumor after TAMg-sEV injection. **(A)** GL261 cells were co-injected with primary microglia-derived sEVs in C57BL/6 mice. Brains were collected at day 7 and tumor presence and size were confirmed by IVIS imaging. **(B)** GL261 cells were co-injected with primary microglia-derived sEVs in C57BL/6 mice. Brains were collected at day 7 and tumors processed for scRNA analysis. tSNE visualization of transcriptional clustering of CD45+ cells infiltrating GL261 glioma in C57BL/6 mice. scRNA-seq analysis identified 18 distinct clusters assigned to 8 major immune cell subtypes. Some clusters contained cells predicted to be of different subtypes; cluster names were assigned based on the most abundant immune cell type. **(C)** Prevalence cells from different subgroups, defined by cluster, cell type, or cluster/cell type. The Y-axis demonstrates the prevalence (% of cells vs total) of a subgroup in TAMg (mice co-injected with primary activated microglia-derived sEVs). The X-axis shows the ratio of prevalence in the TAMg vs. control and Resting-sEVs conditions. The most abundant cell subgroups enriched in TAMg by at least 2-fold compared to other conditions are highlighted in red. **(D)** Heatmap of the top 20 genes upregulated in TAMg-injected macrophages from the most abundant macrophage cluster MF.1. Z-score values for gene expression across TAMg versus control and Resting-sEVs conditions are shown. Perc/pct = percent of cells with gene expression detected with at least 1 read. Magenta = genes upregulated in immune suppressive macrophages. (E) Schematic representation of macrophages differentiation with or without sEVs in vitro. **(F)** Representative flow cytometry graphs of macrophages differentiated in the presence of sEVs isolated from resting or activated (CT2a and GL261) BV2 cells. n=3. Source of macrophages, both male and female mice. (**G-H**) Quantification of the mean fluorescence (MF) intensity of ARG1 (F) and CD206 (G) in macrophages differentiated with sEVs. sEVs isolated from primary resting and activated microglia. One-way ANOVA with Tukey’s multiple comparisons test. n=4. Source of macrophages, both male and female mice. (**I**) T cell suppression assay. Macrophages are collected at day 8 (schematic in D) and co-cultured at a ration 1:4 and 1:8 with T cells. Quantifications of the percentage of proliferating CD8 T cells (normalized to the positive control). sEVs isolated from resting or activated BV2 cells. p-value = Two-way ANOVA with Šídák’s multiple comparisons test. n=3. Source of macrophages, both male and female mice.

Comparative analysis revealed that TAMg-derived sEVs preferentially increased the abundance of monocyte and macrophage populations, which represented the most significantly enriched immune cell subsets compared with tumors receiving microglia-derived sEVs or vehicle control (**Figure 2C** and **Supplementary Table 3**). Because these populations represented predominant responders, we next examined transcriptional changes within the myeloid compartment. Macrophages from TAMg-sEV treated tumors displayed increased expression of genes associated with a pro-tumorigenic phenotype, including the immunosuppressive marker Arg1 (**Figure 2D** and **Supplementary Figure 4C**). Pathway enrichment analysis further revealed activation of pathways associated with neddylation, hypoxia, and cellular metabolism (**Supplementary Figure 4E**), consistent with acquisition of an immunosuppressive macrophage phenotype^29–32^.

To determine whether TAMg-derived sEVs directly promote macrophage differentiation toward an immunosuppressive phenotype, bone marrow-derived monocytes were differentiated *in vitro* in the presence of sEVs isolated from activated or resting microglia (**Figure 2E**). Consistent with our single-cell RNA sequencing data, macrophages differentiated in the presence of TAMg-derived sEVs exhibited increased Arg1 gene expression (**Supplementary Figure 4F**), elevated ARG1 and CD206 protein expression (**Figure 2F-H**), and enhanced suppression of CD8+ T-cell proliferation (**Figure 2I**). Taken together, these findings demonstrate that TAMg-derived sEVs promote the recruitment of bone marrow-derived monocytes and their differentiation into immunosuppressive macrophages within the GBM TME.

### Activated microglia-derived sEVs promote the recruitment of bone marrow-derived myeloid cells into the GBM TME

To validate the immune remodeling identified by single-cell RNA sequencing, we next quantified immune cell infiltration by flow cytometry using both GL261 and CT2a glioma models. Tumors were collected seven days after intracranial co-injection with glioma cells and microglia-derived sEVs (**Figure 2A** and **Figure 3B**). Consistent with our scRNA-seq and immunohistochemical analyses, TAMg-derived sEVs markedly increased the infiltration of CD45+ immune cells into tumors in both glioma models, whereas resting microglia-derived sEVs produced only a modest effect (**Figure 3A and C**).

**Figure 3:**
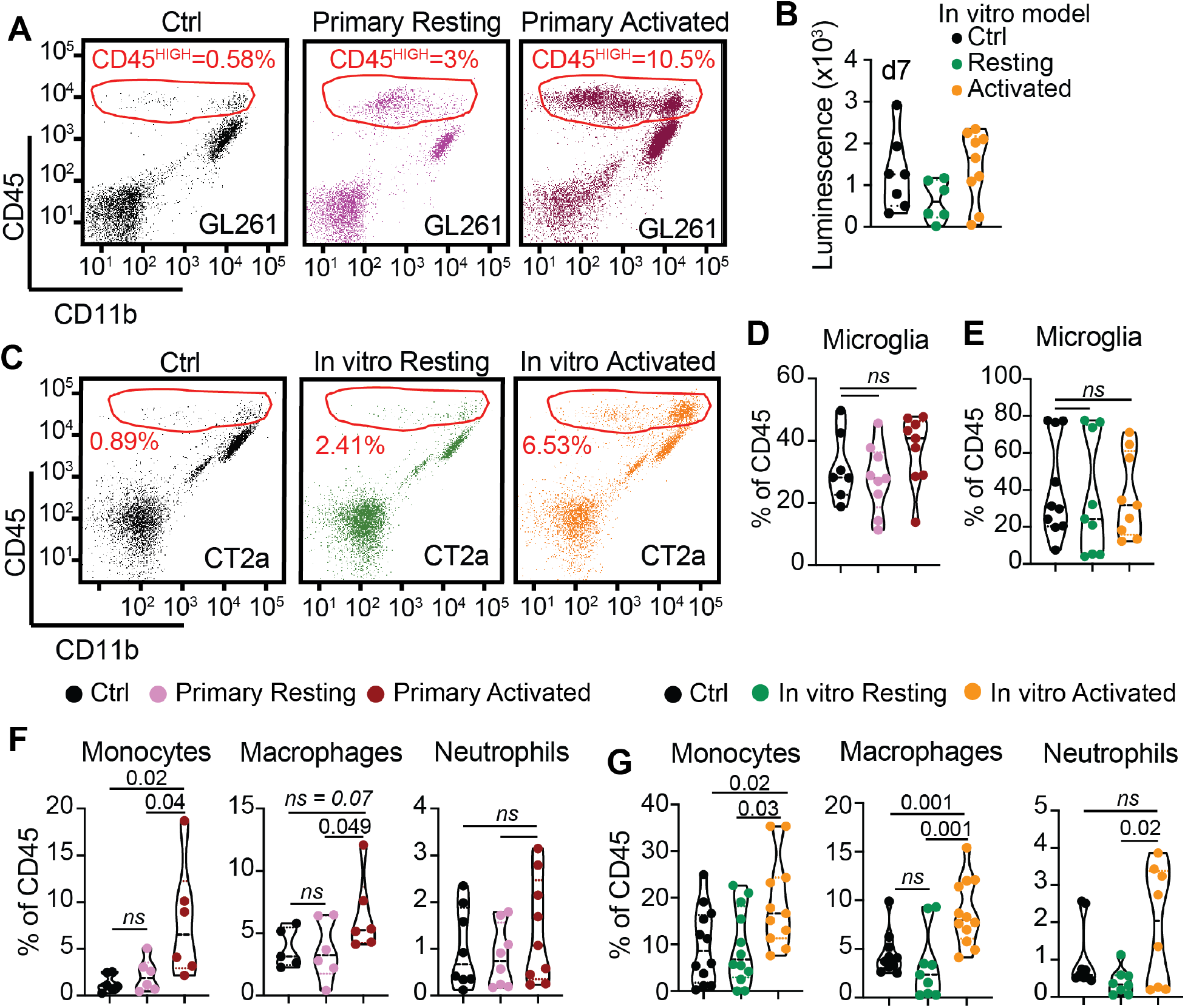
TAMg-sEVs enhance the infiltration of BM-derived myeloid cells into the tumor. **(A)** GL261 cells were co-injected with primary microglia-derived sEVs in C57BL/6 mice. Brains were collected at day 7 and the tumor immune infiltrate was evaluated by flow cytometry. Representative flow panels. CD45^HIGH^ cells = infiltrating immune cells. **(B)** CT2a cells were co-injected with in vitro microglia-derived sEVs in C57BL/6 mice. Brains were collected at day 7 and tumor presence and size were confirmed by IVIS imaging. **(C)** The conditions are in (B) and the tumor immune infiltrate was evaluated by flow cytometry. Representative flow panels. CD45^HIGH^ cells = infiltrating immune cells. **(D)** The conditions are as in (A), and changes in resident microglia were evaluated by flow cytometry after co-injection of GL261 and primary microglia-derived sEVs. Cells were gated in single cells and live cells. n = 6. p-value = One-way ANOVA with Dunnett’s multiple comparison test. **(E)** The conditions are as in (C), and changes in resident microglia were evaluated by flow cytometry after co-injection of CT2a- and BV2-derived sEVs. Cells were gated in single cells and live cells. n = 6. p-value = One-way ANOVA with Dunnett’s multiple comparison test. **(F)** The conditions are as in (A), and changes in infiltrating monocytes, macrophages and neutrophils were evaluated by flow cytometry after co-injection of GL261 and primary microglia-derived sEVs. Cells were gated in single cells and live cells. n = 6. p-value = One-way ANOVA with Dunnett’s multiple comparison test. **(G)** The conditions are as in (C), and changes in infiltrating monocytes, macrophages and neutrophils were evaluated by flow cytometry after co-injection of CT2a- and BV2-derived sEVs. Cells were gated in single cells and live cells. n= 10. p-value = One-way ANOVA with Dunnett’s multiple comparison test.

To determine which immune populations were responsible for this increase, we performed flow cytometric analysis using the gating strategy shown in Supplementary Figure 5A. We found that TAMg-derived sEVs did not alter the proportion of resident microglia within the tumor (**Figure 3D-E**). In contrast, both sources of TAMg-derived sEVs significantly increased the infiltration of bone marrow-derived monocytes (**Figure 3F-G**), consistent with our single-cells RNAseq and immunohistochemical findings. TAMg-derived sEVs also increased macrophage abundance, although this effect was more pronounced following co-injection with *in vitro* activated BV2-derived sEVs (**Figure 3F-G**). By comparison, the proportion of neutrophils, T cells, and B cells was not significantly altered (**Figure 3F-G** and **Supplementary Figure 5B-C**). Together, these findings demonstrate that TAMg-derived sEVs selectively promote the recruitment of peripheral myeloid cells into the GBM TME, supporting the hypothesis that these vesicles drive tumor progression by enhancing bone marrow-derived monocyte infiltration.

### TAMg-derived sEVs migrate to the bone marrow and selectively target monocytes and monocyte progenitors

To investigate how TAMg-derived sEVs promote the recruitment of bone marrow-derived monocytes, we next asked whether these vesicles traffic to the bone marrow and are internalized by resident immune cells. To track their biodistribution *in vivo*, sEVs were labeled with the lipophilic dye DiD and co-injected intracranially with GL261 cells (**Figure 4A**). Bone marrow was collected three days later and analyzed by flow cytometry. DiD-labeled microglia-derived sEVs were readily detected within bone marrow immune cells (**Figure 4B**). Notably, TAMg-derived sEVs were preferentially internalized by monocytes compared with all other immune populations examined (**Figure 4B-C**), whereas resting microglia-derived sEVs showed a weaker and less consistent pattern of uptake. This selective targeting of monocytes, but not neutrophils, was consistent with our previous observation that TAMg-derived sEVs selectively increased monocyte, but not neutrophil, infiltration into GBM tumors (**Figure 3F and I; Supplementary Figure 5B-C**).

**Figure 4:**
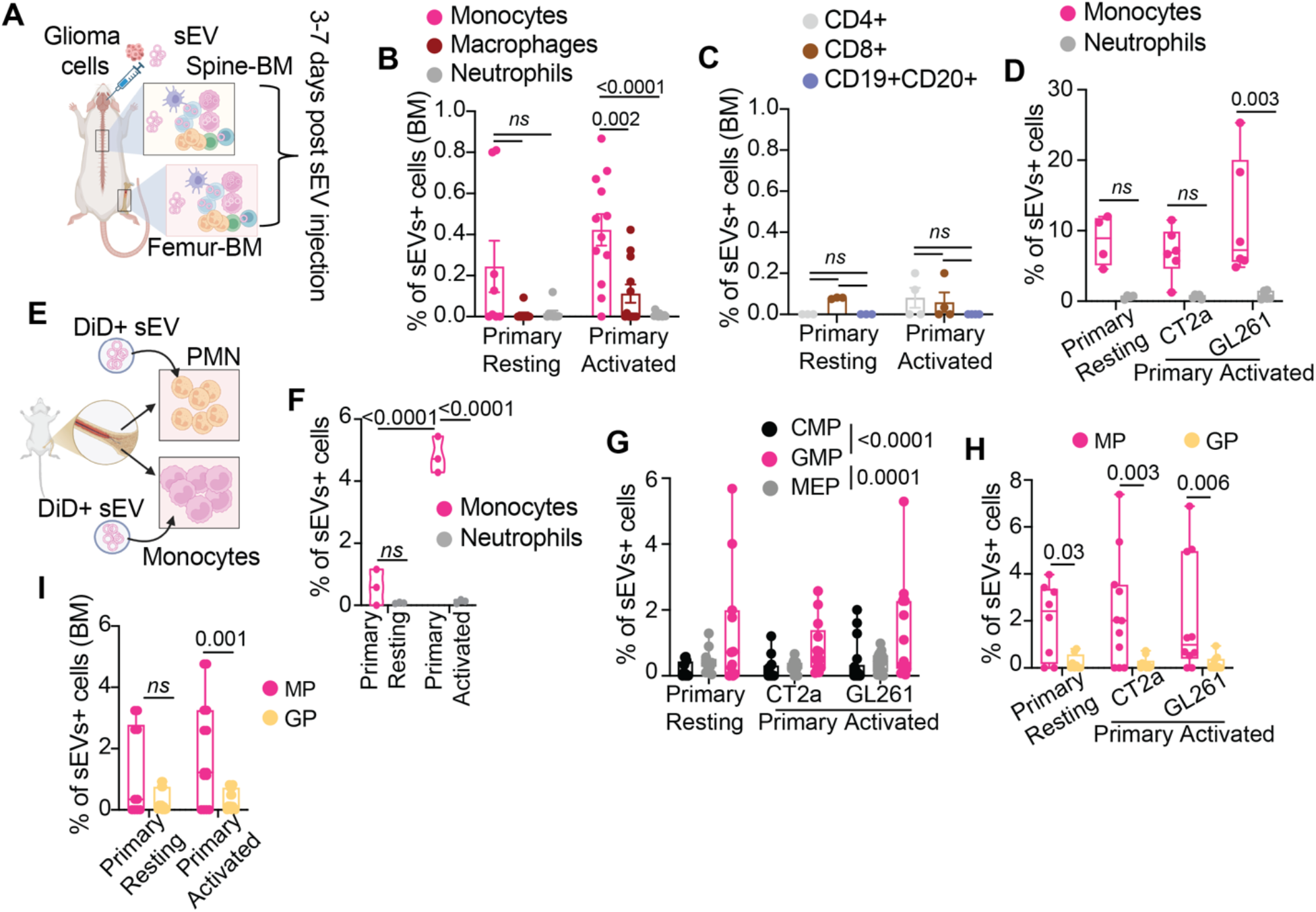
Microglia-derived sEVs are specifically internalized in monocytes and monocyte progenitors. **(A)** Schematic representation of the *in vivo* experiment workflow. Primary microglia-derived sEVs were co-injected with GL261 in 8-week-old naïve female mice and sEVs internalization in immune cell populations in the bone marrow (BM) (both spine and femur) were evaluated after 3 days by flow cytometry. (**B-C**) The conditions are as in (A). Comparison of sEVs internalization between myeloid cells (B; monocytes, macrophages and neutrophils) and lymphocytes (C; CD4+ T cells, CD8+ T cells and B cells (CD19+CD20+)). n= 6. p-value = Two-way ANOVA with uncorrected Fisher’s LSD. **(D)** Immune cells were isolated from the BM of 8-week-old male and female naïve mice and co-cultured with DiD+ sEVs for 24 hours. Comparison between the percentage of sEVs+ monocytes and neutrophils. n= 4-6, both male and female. p-value = Two-way ANOVA with Šídák’s multiple comparisons test. **(E)** Schematic representation of *ex vivo* internalization experiments. Neutrophils (PMN) and monocytes were isolated from the BM of naïve mice and co-cultured with DiD+ sEVs for 24 hours. **(F)** The conditions are as in (E) and the percentage of sEV+ cells was evaluated by flow cytometry. n= 1 female and 1 male. p-value = Two-way ANOVA with Šídák’s multiple comparisons test. (**G-H**) Immune cells were isolated from the BM of 8-week-old male and female naïve mice and co-cultured with DiD+ sEVs. sEVs internalization in myeloid cell progenitors was evaluated after 4 hours in co-culture. (G) CMP = Common Myeloid Progenitors. GMP = Granulocyte Myeloid Progenitors. MEP = Megakaryocyte-Erythroid Progenitors. n= 3. p-value = Two-way ANOVA with Tukey’s multiple comparisons test. (H) sEVs internalization was evaluated in monocyte progenitors (MP) and granulocyte progenitors (GP). n= 3. p-value = Two-way ANOVA with Šídák’s multiple comparisons test. (**I**) The conditions are as in (A). Primary microglia-derived sEVs were co-injected with GL261 in 8-week-old naïve female mice, and sEVs internalization in myeloid cell progenitors in the BM (spine and femur) was evaluated 3 days after intracranial injection. Comparison of sEVs internalization between monocyte progenitors (MP) vs granulocyte progenitors (GP). n= 6. p-value = Two-way ANOVA with uncorrected Fisher’s LSD.

To validate these findings, we performed complementary *ex vivo* uptake assays using bone marrow cells isolated from tumor-free mice. Following 24 hours of co-culture with DiD-labeled sEVs, monocytes efficiently internalized microglia-derived sEVs, whereas neutrophils displayed minimal uptake (**Figure 4D and Supplementary Figure 5D**). To exclude the possibility that this selectivity results from competition between cell populations, purified monocytes and neutrophils isolated from the same mice were cultured separately with identical numbers of DiD-labeled sEVs (**Figure 4E**). Under these conditions, robust sEV uptake was observed exclusively in monocytes, while neutrophils again failed to internalize microglia-derived sEVs (**Figure 4F**), demonstrating that monocytes are intrinsically more permissive to sEV uptake.

We next sought to determine whether TAMg-derived sEVs also target myeloid progenitor populations within the bone marrow (gating strategy shown in **Supplementary Figure 5E**). Among the major progenitor populations, microglia-derived sEVs were preferentially internalized by granulocyte/macrophage progenitors (GMPs; Lin^-^, Sca1^-^cKit^+^, Slam^-^CD34^+^) compared with common myeloid populations (CMPs; Lin^-^, Sca1^-^cKit^+^, Slam^+^CD34^+^) as well as megakaryocyte/erythroid progenitors (MEPs; Lin^-^, Sca1^-^cKit^+^, Slam^-^CD34^-^), irrespective of the sources of the sEVs (**Figure 4G** and **Supplementary Figure 5F**). Because GMPs generate both monocyte and granulocyte lineages, we next compared uptake between monocyte progenitors (MPs; Lin^-^, Sca1^-^cKit^+^, Slam^-^CD34^+^, CD115^+^Ly6C^+^) and granulocyte progenitors (GPs; Lin^-^, Sca1^-^ cKit^+^, Slam^-^CD34^+^, CD115^-^Ly6C^+^). Strikingly, microglia-derived sEVs were selectively internalized by monocyte progenitors, whereas granulocyte progenitors exhibited little to no uptake following four hours of co-culture (**Figure 4H** and **Supplementary Figure 5G**). Consistent with our *ex vivo* findings, DiD-positive monocyte progenitors, but not granulocyte progenitors, were detected in the bone marrow of mice following intracranial administration of DiD-labeled sEVs (**Figure 4I**). Together, these findings demonstrate that TAMg-derived sEVs traffic from the GBM TME to the bone marrow, where they selectively target mature monocytes and monocyte progenitors. Additionally, these data provide a mechanistic basis for the enhanced recruitment of bone marrow-derived monocytes observed following exposure to TAMg-derived sEVs.

### TAMg-derived sEVs induce the expansion of inflammatory and migratory monocytes

We next investigated how TAMg-derived sEVs functionally alter recipient cells following uptake. To test this, CD45+ bone marrow cells isolated from naïve C57BL/6 mice were cultured *ex vivo* in the presence of TAMg-derived or resting microglia-derived sEVs (**Figure 5A**). Flow cytometric analysis revealed that TAMg-derived sEVs selectively increased both the percentage and absolute number of monocytes (CD11b^+^, F4/80^+^, CD115^+^Ly6C^+^) compared with resting microglia-derived sEVs (**Figure 5B-C** and **Supplementary Figure 6C**). In contrast, macrophage and neutrophil populations were not significantly altered (**Supplementary Figure 6A-B** and **D-E**), demonstrating that TAMg-derived sEVs selectively expand the monocyte lineage. To determine whether this expansion resulted from increased proliferation, monocytes were labeled with CellTrace dye and cultured in the presence of sEVs. Following six days of culture, monocytes exposed to TAMg-derived sEVs exhibited significantly greater proliferation than control cells (**Figure 5D**), indicating that TAMg-derived sEVs promote monocyte expansion through enhanced proliferative capacity.

**Figure 5:**
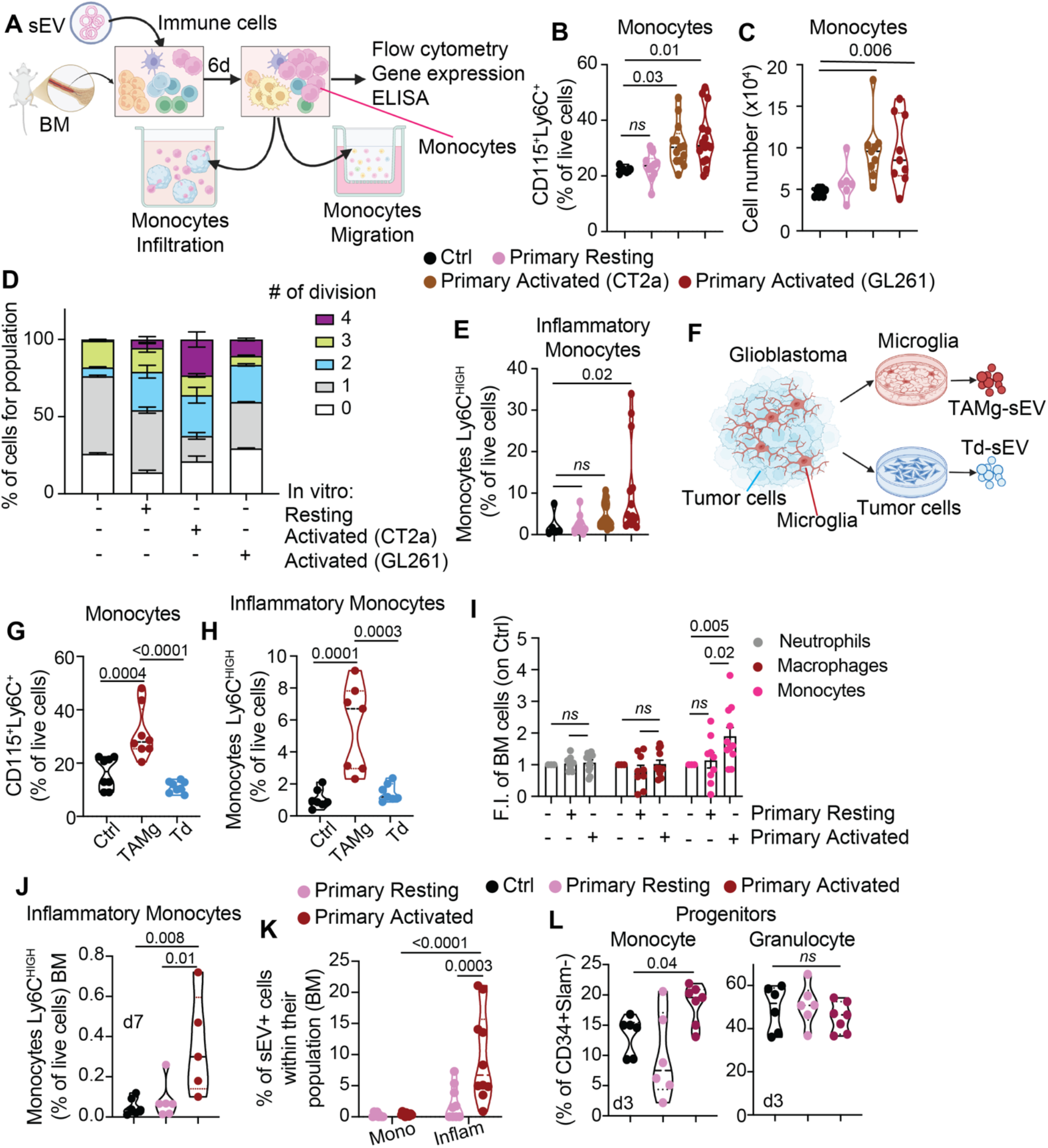
TAMg-sEVs specifically induce an expansion of highly motile inflammatory monocytes, in the bone marrow. **(A)** Schematic representation of experimental workflow. (**B-C**) Cells were isolated from the bone marrow of 8-week-old male and female naïve mice and co-cultured with sEVs. 6 days after seeding, the percentage of monocytes in live cells (B) and the absolute number (C) of monocytes was evaluated by flow cytometry. n = 6. p-value = One-way ANOVA with Kruskal-Wallis test. **(D)** Cells were isolated from the bone marrow of 8-week-old male and female naïve mice, stained with cell trace and co-cultured with sEVs. 6 days after seeding the number of cell division was evaluated by flow cytometry. 1 division: Ctrl vs Activated (CT2a) = 0.03; Ctrl vs Activated (GL261) = 0.01. 2 divisions: Ctrl vs Activated (CT2a) = *ns*; Ctrl vs Activated (GL261) = 0.009. 3 divisions: Ctrl vs Activated (CT2a) = *ns*; Ctrl vs Activated (GL261) = 0.004. n=3. p-value = Two-way ANOVA with Tukey’s multiple comparison test. **(E)** The conditions are as in (B) and the percentage of inflammatory monocytes (Ly6C^HIGH^) was evaluated by flow cytometry. n = 6. p-value = One-way ANOVA with Kruskal-Wallis test. **(F)** Schematic representation of the protocol for isolation of tumor cells and tumor-associated microglia (TAMg) derived sEVs. Tumor cells and microglia were isolated from primary murine tumors. Td = Tumor-derived sEVs. (**G-H**) The conditions are as in (B), and cells were co-cultured with tumor-derived (td) or microglia-derived (TAMg) sEVs. The percentage of monocytes (G) and inflammatory monocytes (H) was evaluated after 6 days. n=3 male and 3 female. p-value = One-way ANOVA with Kruskal-Wallis test. **(I)** C57BL/6 mice were co-injected with GL261 and microglia-derived sEVs (as in Figure 4A). The percentage of neutrophils, macrophages and monocytes in the bone marrow (BM) was evaluated 7 days after the injection. Numbers were normalized on the percentage in control mice. n=9. p-value = Two-way ANOVA with Dunnett’s multiple comparisons test. **(J)** The conditions are as in (I) and the percentage of inflammatory monocytes in the femur bone marrow was evaluated by flow cytometry 7 days post-injection. n=5. p-value = Dunnett’s multiple comparisons test. **(K)** The conditions are as in (I) and DiD+ sEVs were injected. The percentage of sEVs positive monocytes (mono) and inflammatory monocytes (inflam) was measured by flow cytometry three days after the injection. n= 6. p-value Two-way ANOVA with Šídák’s multiple comparisons test. **(L)** The conditions are as in (I) and the percentage of monocyte progenitors and granulocyte progenitors was evaluated by flow cytometry after 3 days of injection in both the spine and femoral BM. n= 6. One-way ANOVA with Dunnett’s multiple comparisons test.

Because inflammatory Ly6C^HIGH^ monocytes represent the major circulating subset recruited to sites of inflammation and tumors^33^, we next examined whether TAMg-derived sEVs preferentially expanded this population. Remarkably, TAMg-derived sEVs selectively increased the proportion of inflammatory monocytes (CD11b^+^, F4/80^+^, CD115^+^, Ly6C^HIGH^; **Figure 5E** and **Supplementary Figure 6F**), suggesting that these vesicles preferentially promote the expansion of the monocyte subset responsible for tumor infiltration. To determine whether this effect was specific to microglia-derived sEVs, we isolated tumor cells and primary microglia from the same GBM-bearing mice (**Figure 5F**) and compared the activity of microglia-derived (TAMg-sEVs) and tumor-derived (Td-sEVs) vesicles. Whereas TAMg-derived sEVs robustly expanded both total and inflammatory monocytes, tumor-derived sEVs failed to induce either response (**Figure 5G-H**), demonstrating that monocyte expansion is a specific property of microglia-derived sEVs.

We next determined whether TAMg-derived sEVs similarly regulate monocyte expansion *in vivo*. To test this, GL261 cells were co-injected intracranially with microglia-derived sEVs, and bone marrow cells were isolated from the spine and femur seven days later (**Figure 4A**). Consistent with our *ex vivo* findings, TAMg-derived sEVs selectively increased monocyte abundance without altering macrophage or neutrophil populations (**Figure 5I** and **Supplementary Table 6**). Furthermore, TAMg-derived sEVs significantly increased the number of inflammatory Ly6C^HIGH^ monocytes (**Figure 5J**). In concordance with the biodistribution studies (**Figure 4**), inflammatory monocytes exhibited the highest frequency of DiD-positive cells following intracranial administration of TAMg-derived sEVs (**Figure 5K**), indicating that this highly migratory subset preferentially internalized microglia-derived sEVs *in vivo*.

Lastly, we investigated whether TAMg-derived sEVs also influence early stages of myelopoiesis. Three days following intracranial administration, TAMg-derived sEVs significantly expanded monocyte progenitors (Lin^-^, Sca1^-^cKit^+^, Slam^-^CD34^+^, CD115^+^Ly6C^+^) without affecting granulocyte progenitors, common myeloid progenitors, megakaryocyte/erythroid progenitors, or granulocyte/macrophage progenitors (**Figure 5L** and **Supplementary Figure 6G**). Together, these findings suggest that TAMg-derived sEVs selectively target the monocyte lineage, promoting expansion of inflammatory monocytes while simultaneously increasing monocyte progenitors within the bone marrow. Notably, these results also suggest that TAMg-derived sEVs remodel bone marrow myelopoiesis to sustain the continuous generation of tumor-promoting monocytes.

### Monocytes expanded by TAMg-derived sEVs acquire an inflammatory and highly migratory phenotype

We evaluated whether monocytes expanded by TAMg-derived sEVs acquire inflammatory and migratory features associated with tumor recruitment. Gene expression analysis of bone marrow myeloid cells cultured for six days with TAMg-derived sEVs revealed marked upregulation of inflammatory genes, including *IL6*, *IL10*, *TNFA*, *IL1B*, and *COX2* (**Figure 6A** and **Supplementary Figure 7A**). In contrast, tumor-derived sEVs failed to induce comparable transcriptional responses (**Supplementary Figure 7B**). Consistent with these findings, TAMg-derived sEVs increased secretion of IL-6 and IL-10 (**Figure 6B**), as well as IL-1β (**Supplementary Figure 7C**). A similar inflammatory gene expression profile was observed in monocytes isolated from the bone marrow of mice co-injected with GL261 cells and TAMg-derived sEVs (**Figure 6C**), confirming that TAMg-derived sEVs promote an inflammatory monocyte phenotype both *ex vivo* and *in vivo*.

**Figure 6:**
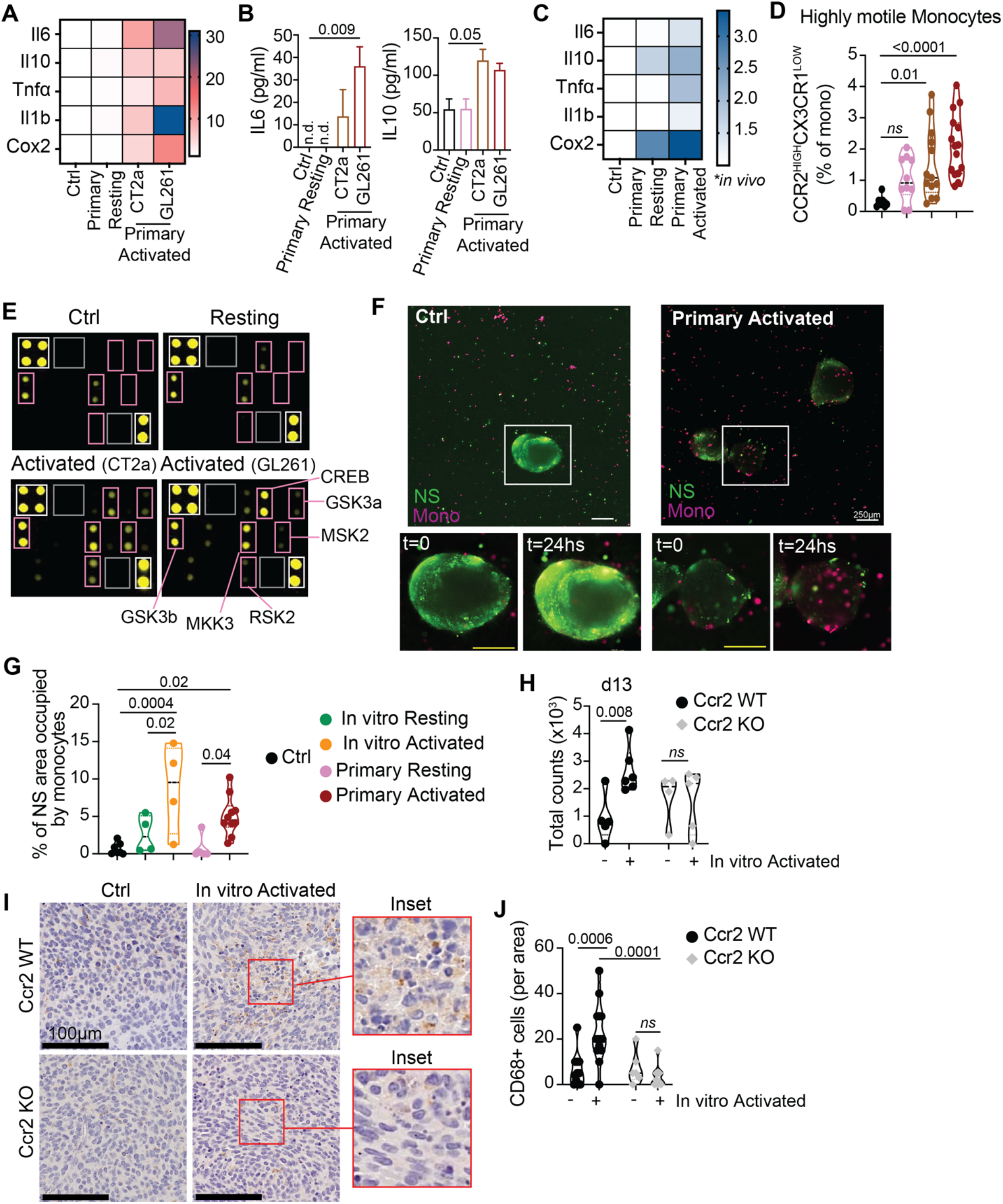
TAMg-sEVs expand highly motile monocytes, which infiltration sustain tumor growth. **(A)** Bone marrow-derived immune cells were co-cultured with sEVs for 6 days. Changes in gene expression were evaluated by qRT-PCR, and values were normalized on control. n=3 females and 3 males. Source of sEVs: primary microglia. **(B)** The conditions are as in (A), and cytokines released in the supernatant was evaluated by ELISA. n=2 males and 2 females. p-value = One-way ANOVA with Dunn’s multiple comparisons test. **(C)** sEVs are co-injected with GL261 cells in 8-week-old C57BL/6 mice and 3 days after the injection, changes in gene expression are evaluated in monocytes isolated from the bone marrow. **(D)** Cells were isolated from the bone marrow of 8-week-old male and female naïve mice and co-cultured with sEVs. 6 days after seeding, the percentage of motile monocytes was evaluated by flow cytometry. n = 6. p-value = One-way ANOVA with Kruskal-Wallis test. **(E)** Bone marrow-derived immune cells were co-cultured with BV2-derived sEVs for 6 days. At day 6 monocytes protein were blotted on a MAPK phosphor Array. White rectangles = positive controls. Gray rectangles = negative controls. Pink rectangles = phospho proteins upregulated by TAMg-sEVs. GSK3b (pT180/Y182). MKK3 (pS189). RSK2 (pS386). CREB (pS133). MSK2 (pS360). GSK3a (pS21). (**F-G**) Neurospheres (NS) generated from tumor-bearing mice (green) were co-cultured for 24 hours with monocytes conditioned with microglia-derived sEVs (magenta). (E) Representative images at different time points of monocytes infiltrating neurospheres. Complete videos are in Suppl. Video 1 and 2. Scale bar = 200μm. White rectangle = inset. (F) Quantification of % of neuropsheres area covered by monocytes. n=3. One-way ANOVA with Dunnett’s multiple comparisons test. (**H**) C57BL/6 WT (Ccr2 WT) and Ccr2-deficient (Ccr2 KO) female mice were co-injected with CT2a cells and BV2-derived sEVs. Brains were collected at day 13 and tumor signal was detected by an IVIS SpectrumCT In Vivo Imaging System. Quantification. n=5. Two-way ANOVA with Dunnett’s multiple comparisons test. (**I-J**) C57BL/6 WT (Ccr2 WT) and Ccr2-deficient (Ccr2 KO) female mice were co-injected with CT2a cell line and BV2 derived sEVs. Brains were collected at day 13 and processed by immunohistochemistry. (I) Representative images of CD68 staining. Objective = 20X. (J) Quantification of CD68+ cells per 10X area. n= 4. Two-way ANOVA with Turkey’s multiple comparison test.

Because inflammatory monocytes are mobilized from the bone marrow and recruited to tumors^8,33,34^ through CCR2-dependent trafficking^35^, we next asked whether TAMg-derived sEVs also induce a migratory phenotype. TAMg-derived sEVs significantly expanded CCR2^HIGH^ monocytes both *ex vivo* (**Figure 6D** and **Supplementary Figure 7D**), and *in vivo* (**Supplementary Figure 7E-F**), suggesting that these vesicles promote the generation of a monocyte population poised for tumor recruitment. To investigate the signaling pathway underlying this phenotype, we examined MAPK pathway activation in response to TAMg-derived sEVs. We found that TAMg-derived sEVs increased the phosphorylation of multiple MAPK-associated proteins (**Figure 6E and Supplementary Original Western Blot 4**), including CREB^36^, GSK3α/β^37^, MKK3, RSK2, and MSK2^38–40^. These factors have established roles in monocyte survival, proliferation, and migration^41–43^, supporting the premise that TAMg-derived sEVs activated a coordinated signaling program that promotes monocyte expansion and motility.

We next tested whether this molecular phenotype translated into enhanced migratory behavior. Toward this goal, bone marrow immune cells from tumor-free mice were cultured with TAMg-derived or resting microglia-derived sEVs for six days, after which equal numbers of monocytes were isolated and analyzed using functional migration assays. Remarkably, monocytes exposed to TAMg-derived sEVs displayed significantly greater transwell migration than control monocytes or monocytes treated with resting microglia-derived sEVs (**Supplementary Figure 7G**). To assess tumor-directed migration in a three-dimensional context, we co-cultured these cells with glioma-derived neurospheres. Despite using equal numbers of monocytes in each condition, only monocytes exposed to TAMg-derived sEVs efficiently migrated toward and infiltrated the neurospheres (**Figure 6F-G, Supplementary Figure 7H-I,** and **Supplementary Videos 1-2**). These findings demonstrate that TAMg-derived sEVs promote the acquisition of a highly inflammatory and migratory phenotype in monocytes.

Finally, we tested whether CCR2-dependent monocyte recruitment was required for tumor-promoting effects of TAMg-derived sEVs *in vivo*. To test this, CT2a cells were co-injected intracranially with TAMg-derived sEVs into wild-type or Ccr2-deficient mice^44^. Whereas TAMg-derived sEVs accelerated tumor growth in wild-type mice, this effect was abolished in Ccr2-deficient mice (**Figure 6H**). Consistent with this result, TAMg-derived sEVs failed to increase infiltration of CD45+ immune cells or CD68+ macrophages in Ccr2-deficient tumors (**Figure 6I-J and Supplementary Figure J-K**). These data establish that CCR2-dependent recruitment of bone marrow-derived monocytes is required for TAMg-derived sEVs to promote GBM progression and identify TAMg-derived sEVs as critical mediators of communication between the GBM TME and the peripheral immune system (**Figure 7**).

**Figure 7.**
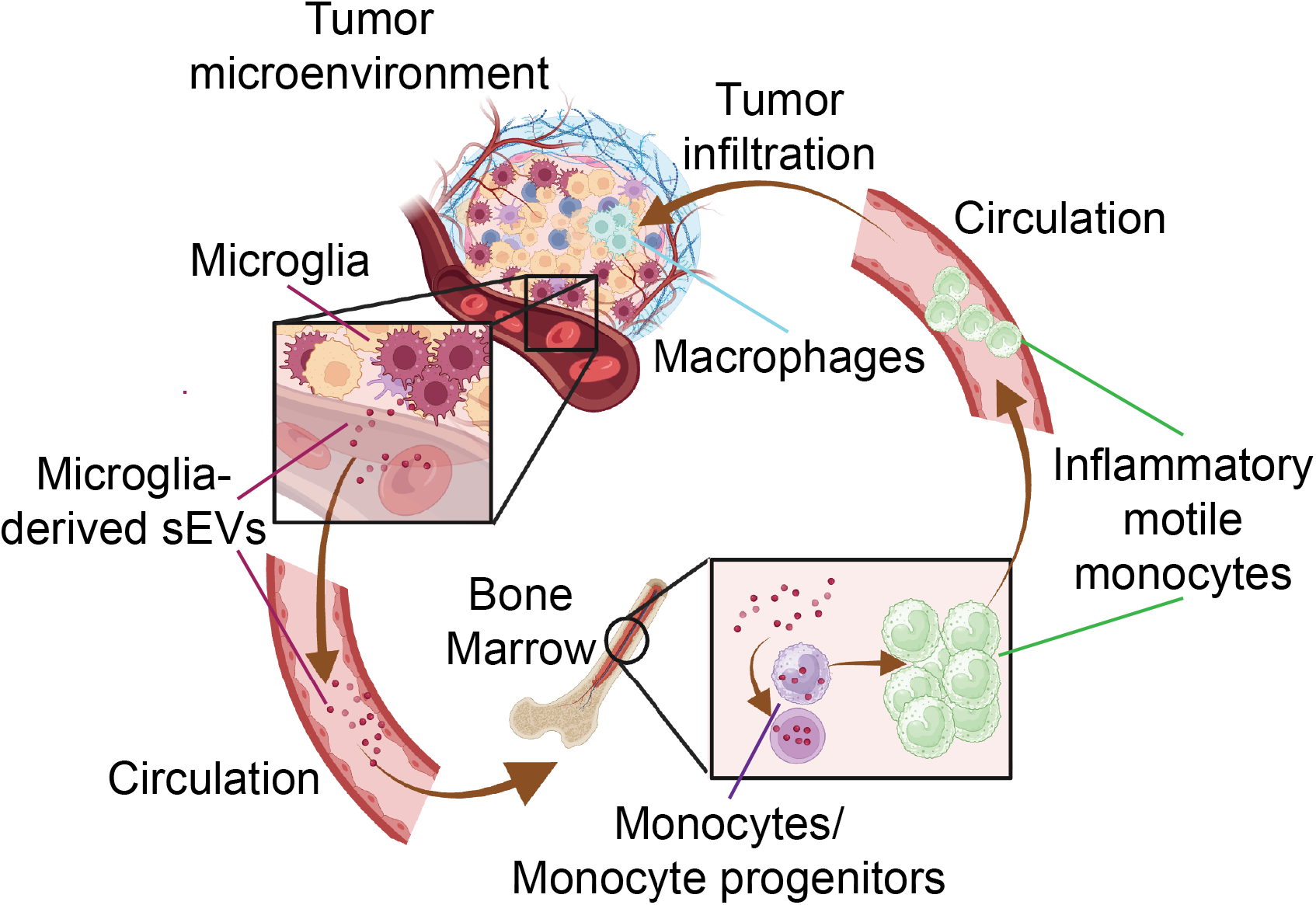
Proposed model highlighting the contribution of TAMg-sEVs on the expansion and mobilization of inflammatory monocytes in glioblastoma.

## DISCUSSION

Glioblastoma (GBM) is characterized by extensive infiltration of peripheral myeloid cells, yet the mechanisms that coordinate their recruitment remain poorly understood. Here, we identify tumor-associated microglia (TAMg)-derived small extracellular vesicles (sEVs) as mediators of communication between the GBM tumor microenvironment (TME) and the bone marrow. Our findings support a model in which TAMg-derived sEVs selectively traffic to the bone marrow, where they are internalized by monocytes and monocyte progenitors, promoting the expansion of inflammatory, highly migratory monocytes that subsequently infiltrate the tumor and support GBM progression. Together, these findings reveal a previously unrecognized mechanism through which resident microglia influence peripheral myeloid cell biology and shape the immunosuppressive TME in GBM.

Microglia have traditionally been viewed as local regulators of GBM progression through the secretion of cytokines, chemokines, and growth factors that promote tumor growth, angiogenesis, and immune suppression^45–53^. Our findings expand this model by demonstrating that tumor-associated microglia also regulate peripheral immunity through extracellular vesicle-mediated communication with the bone marrow. TAMg-derived sEVs selectively targeted mature monocytes and monocyte progenitors, promoted the expansion of inflammatory Ly6C^HIGH^ and CCR21^HIGH^ monocytes, and activated inflammatory transcriptional programs and MAPK-associated signaling pathways linked to proliferation and migration. Functionally, these monocytes displayed enhanced migratory capacity and tumor infiltration, while CCR2 deficiency abolished both increased myeloid infiltration and the tumor-promoting effects of TAMg-derived sEVs. Together, these findings support a model in which tumor-associated microglia influence GBM progression by promoting the generation and recruitment of tumor-promoting monocytes rather than acting solely through local immune modulation.

An intriguing aspect of this response was its remarkable selectivity for the monocyte lineage. Although neutrophils are abundant in both the circulation and bone marrow^33,54^, TAMg-derived sEVs were preferentially internalized by monocytes and monocyte progenitors, whereas neutrophils and granulocyte progenitors exhibited minimal uptake. Consistent with this observation, TAMg-derived sEVs selectively increased monocyte and macrophage infiltration without significantly altering neutrophil abundance. These findings provide one potential explanation for the predominance of monocyte-derived macrophages within the GBM TME. Previous studies have shown that monocyte depletion increases neutrophil infiltration in GBM, suggesting a reciprocal regulation between these myeloid populations^55^. Our findings raise the possibility that selective uptake of microglia-derived sEVs contributes to this imbalance by preferentially expanding the monocyte lineage, although the molecular determinants governing this selectivity remain to be determined.

Collectively, our findings establish tumor-associated microglia as key regulators of systemic myeloid responses through extracellular vesicle-mediated communication with the bone marrow. While additional studies will be required to identify the cargo responsible for monocyte reprogramming and to define how TAMg-derived sEVs cooperate with other signals within the GBM TME, our work identifies a previously unrecognized microglia-bone marrow communication axis that contributes to immune suppression and tumor progression. Targeting the production, trafficking, or uptake of microglia-derived sEVs therefore represents a potential therapeutic strategy to limit pathological myeloid recruitment and improve anti-tumor immunity in GBM.

## MATERIALS AND METHODS

### Mouse strains

C57BL/6J (WT, RRID:IMSR_JAX:000664), B6(Cg)-Tyrc-2J/J (albino, RRID:IMSR_JAX:000058) and B6.129S4-Ccr2tm1Ifc/J (Ccr2 KO, RRID:IMSR_JAX:004999) mice were purchased by The Jackson Laboratory. All experiments were approved by The Wistar Institute Institutional Animal Care and Use Committee.

### Sex as a biological variable

Both male and female mice were used for all *ex vivo* experiments, microglial isolation for extracellular vesicle isolation, in vivo biodistribution experiments, and tumor growth. The sex of the animal used for each experiment is specified in the figure legend. Only female Ccr2 KO mice were used to study differences in tumor growth after co-injection with small extracellular vesicles (sEVs).

### In vivo experiments

For all the *in vivo* experiments, at least 3 animals per group were used. Experiments were repeated at least 2 times. The number of replicates is indicated in the figure legend. The orthotopic murine model was obtained by stereotaxic injection (coordinates: 1.5 mm lateral to the bregma, 0 mm behind, and 3·0 mm ventral to the dura) of tumor cells stably transduced with a luciferase construct with or without sEVs in 2 μL of PBS as described in ^23,24^. For tumor growth, sEVs biodistribution and immune infiltrate experiments, we co-injected 5x10^4^ CT2a-Luc or 10^5^ GL261-Luc with 10^8^ small extracellular vesicles. Tumor growth was followed by IVIS imaging system Living Image Software 4.7.4 (PerkinElmer). Mice were imaged at day 3, 6, 9, 13/14, and 15 days after the injection. The total luciferase counts were normalized on the signal at day 3 and measured as a fold increase over time. or immunohistochemistry analyses, tumors were collected fourteen days after the injection and processed as described below. For studies examining changes in immune infiltrate or scRNA-seq analysis, tumors were collected 7 days after injection. For biodistribution studies, bone marrow was collected three days after the injection.

### Neurospheres generation

To generate tumor-derived neurospheres, 7-week-old C57BL/6 male and female mice were intracranially injected with 5x10^4^ CT2a-Luc. Seven days after the injection, tumors were collected and dissociated as previously described. The negative population from the CD11b+ magnetic bead isolation was collected, washed in DMEM, and seeded in DMEM + 2% sEV-depleted FBS + 20 ng/mL M-CSF. Four days after seeding, tumor cells started generating spheroids. Non-adherent spheroids were collected, transferred to a new plate, and cultured for up to 2 weeks after isolation. At the time of sEVs collection from tumor-derived neurospheres these were maintained in DMEM + 2% FBS-sEVs-depleted FBS for 24 hours. Spheres were cultured for up to 2 weeks after isolation.

### Ex-vivo co-culture experiments with bone marrow-derived immune cells

8-to 10-week-old male and female C57BL/6 tumor-free mice were used. Immune cells were isolated from the bone marrow as described in ^25^. Isolated immune cells were seeded into no-tissue treated 6-well multi-well plates in DMEM 10% FBS + 1 ng/mL M-CSF and incubated with 10^7^ sEVs. After 3 days, fresh medium supplemented with 1 ng/mL M-CSF was added. Six days after seeding, cells were collected for flow cytometry analysis and RNA isolation. Supernatant was collected, centrifuged at 1,000 g x 10 min to remove cell debris, and stored at -80°C for ELISA. As a control, cells were seeded without sEV.

### Ex-vivo macrophages differentiation and T cell suppression assay

8-to 10-week-old male and female C57BL/6 tumor-free mice were used. Immune cells were isolated from the bone marrow as described in ^25^. Isolated immune cells were seeded into no-tissue treated 6-well multi-well plates in DMEM 10% FBS + 10 ng/mL M-CSF to induce monocytes differentiation toward macrophages. At day 3 and 6 medium was replaced with DMEM 10% FBS + 1 ng/mL M-CSF + 10^7^ sEVs. At day 8 differentiated macrophages were collected and characterized by flow cytometry for the expression of ARG1 and CD206 and by qRT-PCR for the expression of Arg1.

For the T cell suppression assay, T cells were isolated from the spleen of - to 10-week-old male and female C57BL/6 tumor-free mice. Splenocytes were labelled with Cell Trace for 10 min at 37 °C and washed twice in complete medium. 2x10^5^ splenocytes were seeded in each well of a U bottom 96-well plate, pre-coated O.N. at 4°C with 1ug/well of anti-CD3 antibody and 0.5ug/well of anti-CD28 antibody. Macrophages collected at day 8 in culture were counted and seeded at different concentration (1 to 4 and 1 to 8, T cells to macrophages). As positive control splenocytes were seeded without macrophages and with CD3 and CD28 stimulation. As negative control splenocytes were seeded without macrophages and stimulation. Three days later cells were collected and stained with a viability dye (Zombie UV), CD3 PE, CD8 BV711 and CD4 APC. The percentage of proliferating cells was measure as dilution of Cell Trace.

### Migration experiments

For 2D migration experiments, six days after seeding, cells were collected, counted and seeded on the upper part of a transwell with 8μm pores in DMEM 10% FBS. DMEM supplemented with 0.5% BSA was placed on the bottom part of the well. The number of migrating cells was counted 2 hours after seeding.

For 3D migration experiments, neurospheres generated as described above were used as in vitro tumor model. Neurospheres were stained with Vybrant™ DiL Cell-Labeling Solution for 20 minutes at 37°C. Excess dye was washed in PBS twice, and spheres were seeded in each well of a µ-Slide 8 Well chamber (Ibidi), previously coated with poly-l-lysine overnight. DiL+ neurospheres were incubated at 37°C overnight to promote attachment to the bottom of the well. Monocytes were collected after 6 days in culture with sEVs, as previously described, stained with Vybrant™ DiD Cell-Labeling Solution for 20 minutes at 37°C, and washed twice in PBS. 10^5^ monocytes were seeded in each well in co-culture with neurospheres. Monocyte movement and migration to the neurospheres were tracked for 24 hours (every 30 min) using a Ti-E inverted microscope (Nikon) equipped with an incubator set at 37°C with 5% CO2 and 95% relative humidity. Digital camera: Hamamatsu ORCA Flash4.0, SN:000641, FOV: 2048 X 2048. Nikon objectives: 10X PLAN FLUOR PH1, N.A. .3, Cal: .653 px\μm, Z-step (slices): 5.6 μm. The Nikon 3D multidimensional file was converted into an extended depth-of-field (EDF) image using NIS-Elements AR, Version: 5.41.02, Build (1711). The files were then exported to the .mp4 format using the H.264 compression codec.

### sEVs internalization experiments

For internalization experiments, isolated sEVs were labeled with the lipophilic dye Vybrant DiD (ThermoFisher, V22887) for 30 min at RT. To remove the unbound dies, after staining, sEVs were processed by size-exclusion chromatography as described above. sEV concentration was measured using ZetaView, and 10 μL aliquots were stored at -80°C for up to 6 months. For *ex vivo* experiments, 10^7^ sEV per condition were used. For *in vivo* experiments, sEVs were concentrated by ultracentrifugation at 100,000 g (2 hours at 4°C) and resuspended to a concentration of 10^8^ sEV/2 μL. Labeled 1X DPBS was used as a control.

For *ex vivo* experiments, 8-to 10-week-old male and female C57BL/6 tumor-free mice were used. Immune cells were isolated as described above. Isolated immune cells were seeded in DMEM 10% FBS into non-tissue-treated 6-well multi-well plates and incubated with DiD+ sEVs for 4 and 24 hours. To evaluate internalization across multiple subpopulations, cells were collected and stained with the myeloid cell progenitor panel, T cell panel, and myeloid cell panel (Supp. Table 2). sEVs internalization was evaluated by flow cytometry.

To evaluate the internalization of sEVs in monocytes or neutrophils separately, we collected the immune cells as described above. Isolated cells were divided into 1 part for neutrophil isolation and 2 parts for monocyte isolation. In both conditions, specific cells were isolated by negative magnetic beads (StemCell, monocytes #19861 and neutrophils #19762). Cells were counted, and 10^5^ cells were incubated with 10^7^ sEV for 4 hours. sEVs internalization and purity of the isolation were evaluated by flow cytometry.

For *in vivo* experiments, 10^8^ DiD+ sEVs were co-injected with 10^5^ GL261-Luc cell line. Three days after the intracranial injection, both the spine (vertebrae) ^26^ and the femur (long bones and ilia) bone marrow were collected, cells isolated, and stained with the myeloid cells and myeloid progenitors flow panels (Suppl. Table 2). sEVs internalization was evaluated as % of DiD+ cells.

### Statistical analyses

Statistical analysis was performed using GraphPad Prism 10 software. All *in vitro* biological experiments were performed at least 3 times with both male and female mice. In vivo experiments were repeated at least 2 times; the number of animals is specified in the figure legends. For comparing 3 or more groups, a One-way ANOVA was used. To determine how a response was affected by 2 factors, a two-way ANOVA test was used. For longitudinal tumor growth, a two-way ANOVA for repeated measures was used. Statistical tests used are specified in the figure legends.

## Supplementary information

This manuscript has 8 supplementary figures, 2 supplementary videos and 8 supplementary tables.

## ACKNOWLEDGMENTS

Small extracellular vesicles characterization was performed with the support of the Extracellular Vesicles core at University of Pennsylvania (RRID: SCR_022444).

## Funding

This work was supported by The W.W. Smith Charitable Trust (I.B.) and R01 CA316572 (T.B.).

## Author contributions

IB conceived the project. IAR and MQP performed experiments on sEVs characterization, modulation of myeloid cell expansion, sEVs internalization experiments, and migration assays. IAR, MQP, MY, and IB characterized the effect of sEVs on tumor growth and immune cell infiltration *in vivo*. NP tested the effect of sEVs on tumor cells in vitro. NP and AM assisted with characterizing the effect of sEVs on monocytes in vitro. TB reviewed and edited the manuscript. QL assisted with the statistical analyses. AVK analyzed bioinformatics data. IAR, MQP, AVK, and IB analyzed the data. IB wrote the manuscript.

## Competing interests

The authors declare that they have no competing interests.

## Data and materials availability

Raw single-cell RNA-seq data were deposited to the NCBI GEO repository under accession number GSE316676.

## Notes

### Competing Interest Statement

The authors have declared no competing interest.

