## Supplementary material for "Microglia-derived Small Extracellular Vesicles Promote Bone Marrow Monocyte Expansion and Recruitment in Glioblastoma"

Supplementary information included: Supplementary methods, 7 supplementary figures and 8 supplementary tables.

### SUPPLEMENTARY METHODS

#### Antibodies and reagents:

Vybrant™ DiD and DiL Cell-Labeling Solution and Fc block were purchased from ThermoFisher. Zombie UV™ Fixable Viability Kit was purchased from Biolegend (#423107). Antibodies to Calnexin (Cat# 2679, RRID:AB_2228381), GM130 (Cat# 12480, RRID:AB_2797933) and β-actin (Cat# 5125, RRID:AB_1903890) were obtained from Cell Signaling. Antibodies to Tsg101 (Cat# 14497-1-AP, RRID:AB_2208090), ALIX (Cat# 12422-1-AP, RRID:AB_2162467), CD9 (Cat# 20597-1-AP, RRID:AB_2878706) and CD81 (Cat# 27855-1-AP, RRID:AB_2880995) were from ProteinTech. Antibody to CD9 (Cat# AHS0902, RRID:AB_2536363) was purchased from Thermo Fisher Scientific.

#### Cell culture:

The murine glioblastoma cell line CT2a-Luc was purchased from Millipore-Sigma (Millipore Cat# SCC194, RRID:CVCL_ZJ44). Murine glioblastoma cell lines GL261-Luc (#ABC-RC084H) and BV2 (ABC-TC212S) were purchased from Accegen. All cell lines were grown in DMEM (Corning) supplemented with 10% fetal bovine serum (FBS) (Gemini) and 1% Penicillin/Streptomycin (Corning) and maintained in a humidified incubator at 37°C with 5% CO2. Cell passaging was limited to fewer than 30 passages. Cultured cells were verified to be mycoplasma-free using the Mycoplasma Primer Set (Bioo Scientific).

#### Primary microglia isolation and ex-vivo culturing for sEVs collection:

Microglia were isolated from the brain of female and male 8-week-old C57BL/6 mice. For Primary Activated microglia either 5 x $10^4$ CT2a-Luc or $10^5$ GL261-Luc cells were intracranially injected in 7-week-old mice as previously described. Successful tumor cell injection was verified by IVIS imaging on days 3 and 6 after injection. 7 days after the injection, brains were harvested and tumor localization was identified using the IVIS imager prior to being resected. Microglia were isolated from eleven C57BL/6 female mice (6 injected with GL261 and 5 injected with CT2a) and nine C57BL/6 male mice (3 injected with GL261 and 6 injected with CT2a). For Primary Resting microglia, brains were collected from 8- to 10-week-old tumor-free mice. Microglia were isolated from five C57BL/6 female mice and four C57BL/6 male mice. Under all conditions, the brain and tumor were dissociated using the Adult Brain Dissociation Kit (Miltenyi, #130-107-677) according to the manufacturer's instructions. Microglia cells were enriched by magnetic bead isolation (StemCell CD11b kit, #18970) and seeded on poly-l-lysine-coated wells in a 6-well multi-well plate in DMEM containing 10% sEV-depleted-FBS, 1% penicillin/streptomycin, 2 mM L-glutamine, and 20 ng/mL M-CSF. After seeding, cells were maintained by replacing 50% of the medium every other day and their supernatant was collected every 24 hours, on days 6, 7, and 8. At the end of the experiment (day 8), microglia cells were collected for flow cytometry analysis, RNA, and protein isolation. The purity of the microglia preparation was evaluated by flow cytometry. Cells were stained with: CD11b, F4/80, CD206, and TMEM119 (Supp. Table 2). Microglia cells were gated in the single cells-live population as CD11b+F4/80+ and TMEM119+ (Suppl. Fig. 1B). CD206 or ARG1 were used as markers of activation.

**BV2 activation for sEVs production:**

To generate the tumor supernatant for BV2 activation, 7-8-week-old C57BL/6 female and male mice were intracranially injected with either 5 x $10^4$ CT2a-Luc or $10^5$ GL261-Luc cells, as previously described. Successful tumor cell injection was verified by IVIS imaging at days 3 and 6 after injection. 7 days after the injection, tumors were collected and maintained in culture for 24 hours in DMEM medium supplemented with 2% sEV-depleted-FBS. The collected supernatant was centrifuged at 1,000 g for 10 min at 4°C to remove cells and debris, then stored at -80°C. For sEV production *in vitro*, BV2 cells were seeded in 10cm dishes, and after 24 hours (when at 60-70% confluence); the medium was replaced with a 1:1 mix of tumor supernatant and empty medium. Cells were activated for 6 hours, then the tumor supernatant-enriched medium was removed, cells were washed once in 1X DPBS, and DMEM supplemented with 2% sEV-depleted FBS was added. 24 hours later, medium was collected for sEV isolation. RNA was isolated after each activation to confirm proper cell activation by qRT-PCR (Supp. Table 1). An equivalent number of BV2 cells was always used as a resting control (In vitro Resting). Cells were seeded and followed the same protocol, but no tumor supernatant was added; cells were activated with normal DMEM supplemented with 2% sEV-depleted FBS. sEV-depleted-FBS was generated by centrifugation at 100,000 g at 4ºC for 18 hours.

**Small extracellular vesicles isolation and characterization:**

Medium was centrifuged immediately after collection at 1,000 g for 10 min at 4°C to remove cell debris. The supernatant was concentrated using Amicon Ultra-15 centrifugal filter tubes (Merk Millipore) at 4,000 g for 40 min at 4°C. Larger extracellular vesicles were eliminated by centrifugation at 10,000 g for 30 min at 4°C. Finally, sEVs were isolated by qEV size-exclusion column (SEC–iZON) as described in [25]. 500 µL aliquots of the concentrated supernatant were loaded onto qEV columns and eluted with PBS. 2mL of sEVs was collected in an Eppendorf tube corresponding to fractions 7-8-9-10. To confirm the presence of sEVs, aliquots of 20 µL of sEVs diluted in 2 mL of 0.2µm filtered water were analyzed using ZetaView NTA (Particle Metrix). The zeta potential of collected sEVs was measured using the ZetaView system. Expression of sEV biomarkers CD63, CD9, and CD81 was quantified using an ExoView R100 system (NanoView Biosciences). After quantification, sEVs were stored at –80°C as single-use aliquots to avoid freeze-thaw cycles, for a period not exceeding 3 months [26].

**Immunoblotting:**

Firstly, sEVs were precipitated by ultracentrifugation at 100,000 g (2 hours at 4°C) and resuspended in 100 µl of RIPA buffer. sEVs protein content was measured by Bradford assay, and 15 µg of protein were used for Western blot analysis. sEVs were denatured in Laemmli buffer at 96°C for 5 min, and the protein extracts were resolved by SDS-PAGE. Antibodies were tested against Alix, β-actin, Calnexin, CD9, CD81, GM130 and Tsg101. Antibodies were chosen according to MISEV 2023 guidelines [18]. SuperSignal™ West Pico PLUS Chemiluminescent Substrate (Thermo Fisher Scientific) was used to visualize the bands. Membranes were imaged, and bands were quantified using the LI-COR ODYSSEY® M.

**Super resolution experiments:**

1.5 Glass Coverslips 12mm Round (Thomas #1217N79) were placed in a 24 -well multiplate and coated with 100 μL of poly-l-lysine overnight at 4°C. The following day, coverslips were washed twice with 1X DPBS and 500ul of in vitro resting or in vitro activated sEVs diluted in 1X DPBS were added to individual coverslips and incubated at 4°C overnight on agitation. Coverslips were then washed twice with 1X DPBS and blocked with 500 μL of 2% BSA in 1X DPBS for 1 hour at room temperature on agitation. Primary antibodies, mouse anti-mouse CD9 and rabbit anti-mouse CD81, were diluted 1:100 in 0.2% BSA in 1X DPBS. On day 4, after 3 washes in 1X DPBS, coverslips were incubated with their respective secondary antibody, either Rabbit IgG (H&L) Antibody ATTO 647N Conjugated (Rockland Cat# 611-156-122, RRID:AB_10893043) or Mouse IgG (H&L) Antibody ATTO 594 Conjugated (Rockland Cat# 610-155-121, RRID:AB_10893162), diluted 1:500 in 1X DPBS for 1 hour at room temperature, protected from light and on agitation. Coverslips were then washed 3 times with 1X DPBS and mounted onto glass slides using Prolong Diamond Antifade (Thermo Fisher Scientific #P36965) mounting media. Slides were stored at 4°C for at least 48 hours before imaging. Images were captured with a Leica Stellaris 8 3X TauSTED scanning laser confocal microscope (Leica Microsystems, Deerfield, Illinois, USA) incorporating a white-light laser, a 775-pulsed depletion laser, and a 100X/1.4 oil-immersion objective. Acquisition parameters were set in Leica LAS-X software to 1024 x 1024 pixels and 5X zoom for a resolution of 22 nm/pixel.

**Flow cytometry analysis:**

For surface staining cells were incubated with an antibody mix for 15 min at room temperature (RT). For intracellular staining, cells were incubated 15 min at RT with BD Cytofix/Cytoperm™ Fixation/Permeabilization Kit (BD, #554714), washed with BD Cytofix/Cytoperm wash buffer and incubated with the antibody mix for 15 min at RT. For intranuclear staining cells were permeabilized with FoxP3 permeabilization buffer (eBioscience, #00-5523-00) for 45 min at 4°C and incubated with the antibody for 45 min at 4°C.

To assess changes in immune infiltration within the tumor, brains were collected seven days after the injection, and the tumor area identified by IVIS imaging was resected. Tumors were dissociated using the Adult Brain Dissociation Kit (Miltenyi, #130-107-677) following manufacturer's instructions. Cells were then resuspended in MACS buffer (1X DPBS, 4mM EDTA and 1% FBS), seeded in a 96-well multiplate and stained as described. Complete list of antibodies in Supplementary Table 7.

**Immunohistochemistry staining:**

Paraffin-embedded tissue sections were deparaffinized and rehydrated through sequential incubations in xylene (or xylene substitute), graded ethanol solutions (100% and 95%), and distilled water. Antigen retrieval was performed by heat-induced epitope retrieval (HIER) in EDTA buffer (Target Retrieval Solution, Dako) at 95 °C for 20 min using a pressure cooker. Slides were then allowed to cool at room temperature for 30 min. Endogenous peroxidase activity was quenched by incubation in 3% hydrogen peroxide for 10 min, followed by washes in PBST. Tissue permeabilization was achieved by incubating with Triton X-100 in PBS for 15 min. Non-specific binding was blocked with 2.5% normal horse serum for 1 h at room temperature. Sections were incubated overnight at 4°C with primary antibodies diluted in Da Vinci Green Diluent (Biocare Medical): anti-CD45 (1:200; Cell Signaling Technology Cat# 70257, RRID:AB_2799780) and anti-CD68 (1:2000; Proteintech Cat# 28058-1-AP, RRID:AB_2881049). After washing, sections were incubated with an HRP-conjugated polymer secondary antibody (ImmPRESS HRP Horse Anti-

Rabbit IgG, Vector Laboratories) for 30 min at RT. Immunoreactivity was visualized using DAB as chromogen, followed by rinsing in running tap water. Sections were counterstained with Harris hematoxylin, blued in lithium carbonate solution, dehydrated through graded ethanol and xylene, and mounted using a solvent-based mounting medium.

**ELISA:**

Proteintech ELISA kits were used to detect mouse IL6 #KE10007, mouse IL10 #KE10103 and mouse IL1β #KE0003. Cell culture supernatant samples were run in duplicate, with 100ul added per well and the plate incubated at 37ºC for 2 hours. Next, 100ul of Detection Antibody 1X solution was added to each well and the plate was incubated for 1 hour at 37ºC. Then 100 µL of 1X Streptavidin-HRP solution was added to every well followed by a 40-minute incubation at 37C. The plate was manually washed 4 times using Wash Buffer 1X after each incubation. 100 µL of TMB substrate solution was added to each well and the plate was incubated for 15 minutes in the dark at 37ºC. Without washing the plate, 100 µL of Stop Solution was added to each well, and the absorbance was measured at 450 nm using a microplate reader.

**RNA isolation and qRT-PCR:**

Cells were lysed using the TRI Reagent® (Zymo Research, R2050-1) and stored at -80°C until processing. Total RNA was extracted using the Direct-zol™ RNA Miniprep kit (Zymo Research) according to the manufacturer's instructions. Complementary DNA (cDNA) was synthesized with the High-Capacity cDNA Reverse Transcription Kit with RNase Inhibitor (Thermo Fisher Scientific). Quantitative real-time PCR was performed using with SYBR™ Green PCR Master Mix for 18S (as the housekeeping gene) and the target genes listed in the Suppl. Table 8.

**MAPK Pathway Phosphorylation Array:**

10-week-old female C57BL/6 tumor-free mice were used. Immune cells were isolated from the bone marrow as described in [27]. Isolated immune cells were seeded into no-tissue treated 6-well multi-well plates in DMEM 10% FBS + 1 ng/ml M-CSF and incubated with $10^7$ sEVs. After 3 days, fresh medium supplemented with 1 ng/ml M-CSF was added. Six days after seeding, cells were collected and proteins isolated. 250μg of proteins were used for each sample. Human and Mouse MAPK Pathway Phosphorylation Array (Cat# AAH-MAPK-1-4, RayBiotech) was used to detect phosphorylated proteins in the samples. Samples were incubated following the manufactorer's instruction and signal was capture using the LI-COR ODYSSEY® M.

**scRNAseq:**

Female 8-week-old C57BL/6 mice were intracranially injected with $10^5$ GL261-Luc cells in 2 μL of PBS (Ctrl), sEV isolated from female primary resting microglia, and sEV isolated from tumor-associated microglia isolated from a GL261-injected tumor. Successful tumor cell injection was verified by IVIS imaging at days 3 and 6 after injection. Seven days after the injection, brains were collected, and the tumor area was identified using the IVIS imager and resected. Tumors were dissociated and stained with Zombie UV for detecting live cells and CD45 to isolate immune cells. CD45+ live cells were isolated by fluorescence activated cell sorting with gating on live single cells using a MoFLoAstrios flow cytometer. After sorting, cells were counted with Trypan blue and a viability higher than 95% was confirmed. 29,000 single-suspension cells were partitioned using the 10x Genomics Chromium iX single-cell platform (10x Genomics, Pleasanton, CA) to capture an expected 20,000 single cells per sample. 3 lanes were loaded in total with 3 different samples. Single-cell droplets were generated using the Chromium GEM-X single-cell 3' kit v3 (10x Genomics). cDNA synthesis and amplification, library preparation, and indexing were done using the 10x Genomics Gene Expression Library Preparation kit (10x Genomics), according to the manufacturer's instructions. The following PCR cycles were used at each step: 11 PCR cycles for cDNA amplification, and 12 PCR cycles for index incorporation into the final gene expression library. Overall library size was assessed using the 4200 TapeStation and the High-sensitivity DNA 5000 screentape (Agilent, Santa Clara, CA). Concentrations were determined using the Qubit Fluorometer 2.0 (Thermofisher, Waltham, MA). Gene expression libraries were pooled according to the reads per cell requirement suggested by 10x Genomics. Next Generation Sequencing on the pool was done on the NextSeq 2000 (Illumina, San Diego, CA) using a P3 100 cycle kit (Illumina), yielding 1.2B total reads, paired-end run with the following run parameters: 28 base pair x 10 base pair (index) x 90 base pair.

The Cell Ranger Software Suite v7.0.1 (https://support.10xgenomics.com) was used to preprocess scRNA-seq data using mm10 transcriptome information using STAR [29]. A common single-cell pipeline will be used to analyze the data using the Seurat v5 package [30] on the output of Cellranger. Doublets identification was performed using R package scDblFinder [31]. The low-quality cells with few expressed genes (n<200) and mitochondrial content of more than 10% were removed from analysis. After filtering, data from different samples were integrated using 2000 anchor genes and clustered using the default resolution of 0.8, and subtypes of infiltrating and pancreatic cells were performed using the SingleR R package [32] subtyping approach with ImmGenData reference. After sample integration, clustering, and immune cell subtyping, cells types with <5% prevalence in the cluster were removed (n=322). Clustering was evaluated visually using t-SNE or UMAP with default parameters. Significance of cluster/cell subtype-specific genes and genes differentially expressed between conditions was estimated using the Wilcoxon rank sum test, and FDR<5% genes were a subject of downstream enrichment analyses. Enrichment analysis was performed using QIAGEN's Ingenuity® Pathway Analysis software (IPA®, QIAGEN Redwood City,www.qiagen.com/ingenuity) using “Canonical pathways” and “Upstream Regulators” options. Pathways and regulators that passed FDR<5% and were predicted to be activated or inhibited based on their activation-state Z-scores were reported. IL1A signature scores were calculated for each cell using the average expression of 26 IL1α-regulated genes as derived from the Ingenuity Knowledgebase. Raw single-cell RNA-seq data were deposited to the NCBI GEO repository under accession number GSE316676.

### SUPPLEMENTARY FIGURES

**Supplementary figure 1. Tumor-associated microglia models.**

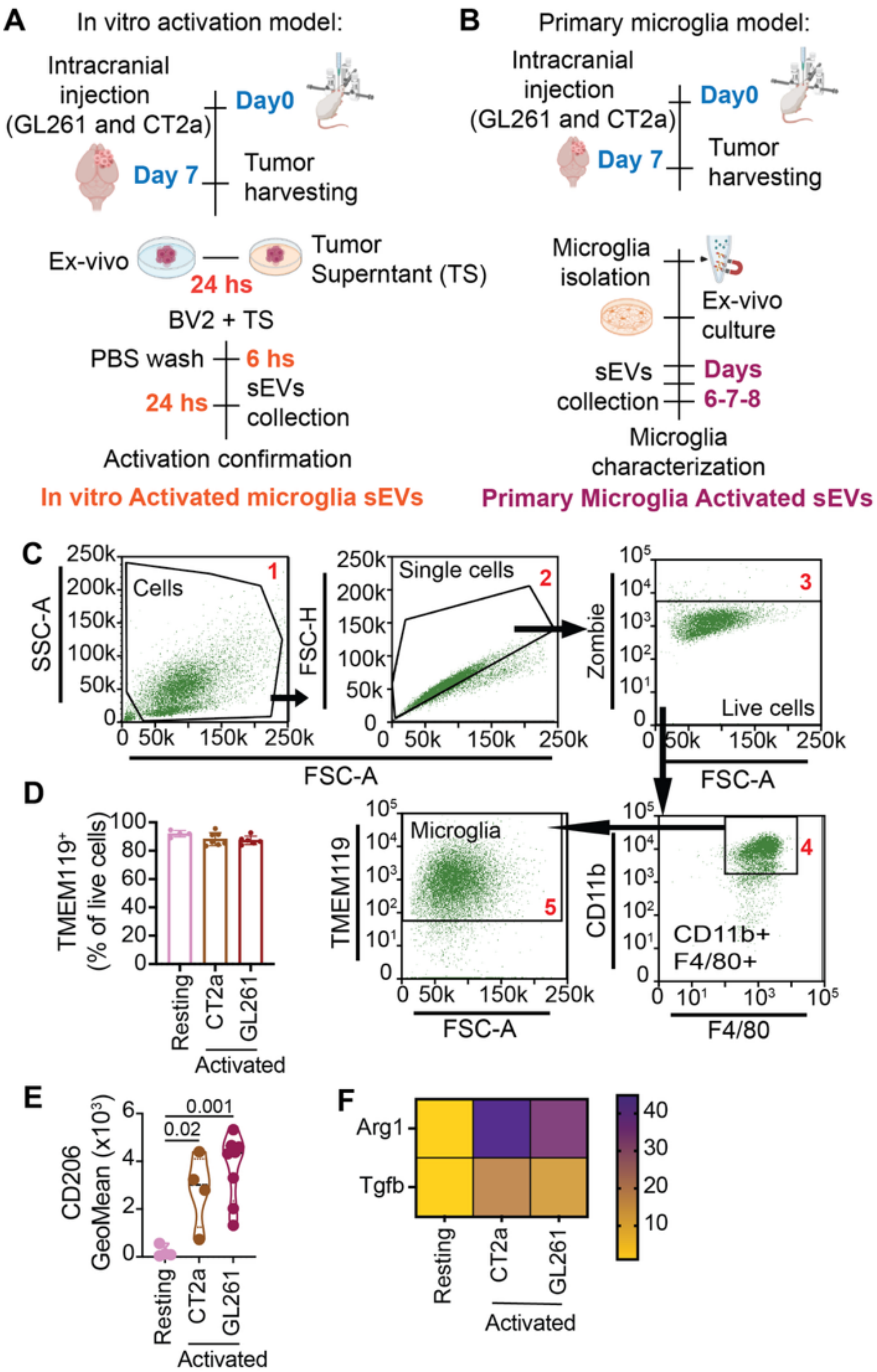


(**A-B**) Schematic workflow of the generation of the microglia models used. (A) In vitro models: BV2 microglia cell line is activated with tumor supernatant (TS) conditioned with tumor isolated from mice injected with CT2a or GL261 cells. (B) Primary model: mice were injected with CT2a or GL261 cells, 7 days after the injection microglia were isolated, maintained in culture for up to 8 days and sEVs collected.

(**C**) Gating strategy for the characterization of primary microglia isolated from naïve or tumor bearing mice. Representative graphs.

(**D**) The purity of primary microglia cells was confirmed by flow cytometry as TMEM119+ cells. One-way ANOVA. Resting: n=4 primary resting microglia. CT2a: n=7 activated microglia isolated from mice injected with CT2a. GL261: n=6 activated microglia isolated from mice injected with GL261.

(**E**) CD206 levels evaluated by flow cytometry in primary microglia isolated from the brain of naïve mice (resting) or tumor bearing mice (injected with CT2a or GL261). One-way ANOVA. Resting microglia: n= 4 (2 males and 2 females). CT2a: n= 4 (2 males and 2 females). GL261: n= 8 (5 females and 3 males).

(**F**) Arg1 and Tgfb expression evaluated by qRT-PCR in primary microglia models, valued normalized on expression levels in primary resting microglia. Resting = microglia isolated from the brain of naïve mice. CT2a: activated microglia isolated from mice injected with CT2a. GL261: activated microglia isolated from mice injected with GL261. n = 3.

**Supplementary figure 2. Small extracellular vesicles (sEVs) characterization.**

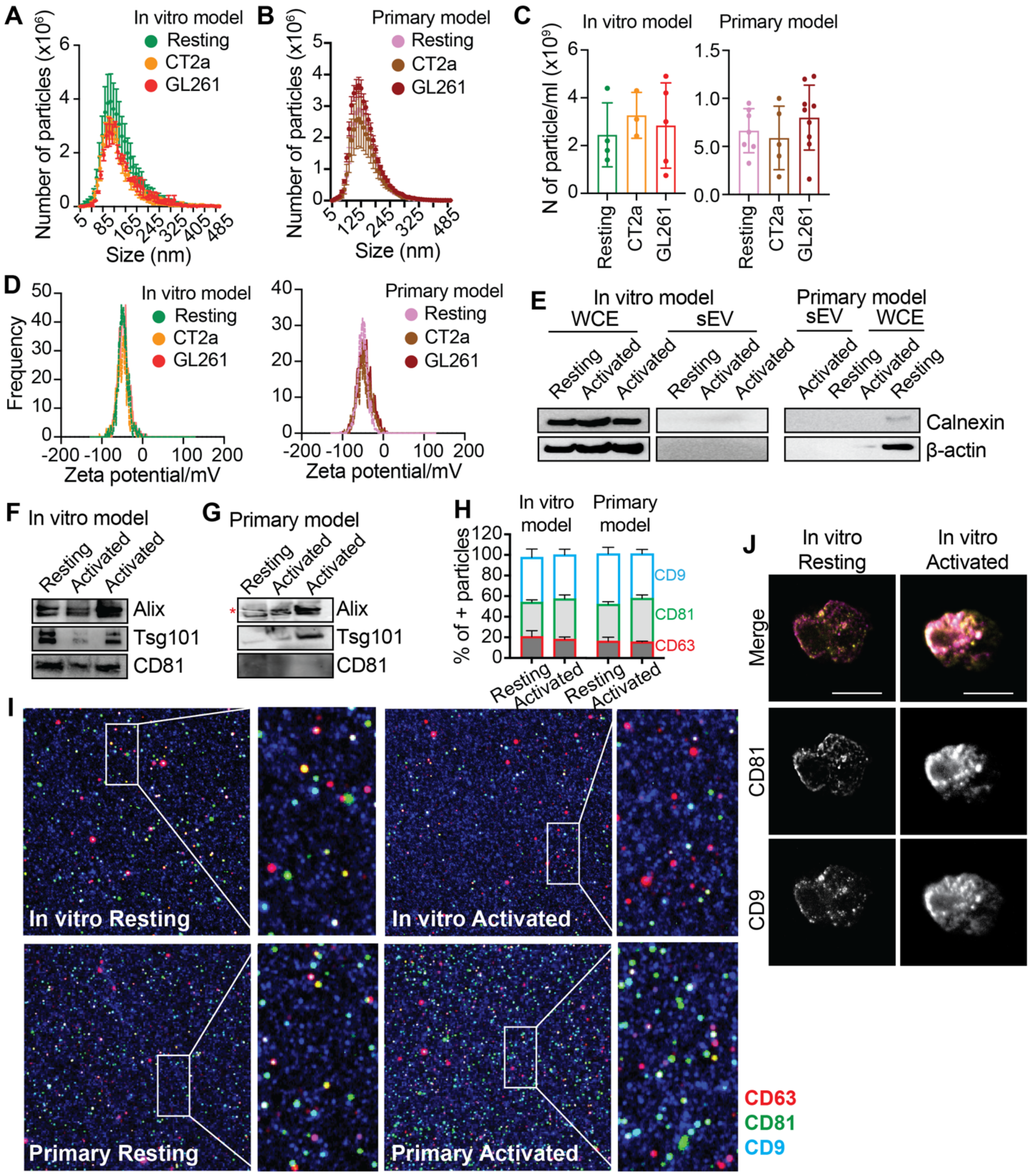


(**A-B**) Size distribution and concentration of isolated sEVs. (A) sEVs isolated from BV2 cells (in vitro model). Resting: non activated BV2. CT2a: BV2 activated with CT2a tumor supernatant. GL261: BV2 activated with GL261 tumor supernatant. (B) sEVs isolated from primary microglia models. Resting: microglia isolated from the brain of naïve mice. CT2a: microglia isolated from CT2a tumor bearing mice. GL261: microglia isolated from GL261 tumor bearing mice. n = 5.

(**C**) Number of particles per ml of supernatant evaluated by ZetaView. Each dot represents one sample.

(**D**) Zeta potential evaluated by ZetaView in sEVs produced by BV2 cell line (left) and primary microglia (right). n = 5.

(**E**) Western blot evaluation for lack of contamination markers in sEVs produced by BV2 cell line (in vitro) and primary microglia. Calnexin = ER marker. β-actin = cellular marker. WCE = whole cell extract.

(**F-G**) Western blot evaluation for the presence of sEV markers in sEVs produced by BV2 cells in vitro (F) and primary microglia models (G). Red asterisk (*) = broken membrane.

(**H-I**) Evaluation of tetraspanin (CD9, CD81 and CD63) expression on sEVs using ExoView system. n = 3. H = quantification. I: Representative ExoView images of sEVs produced by BV2 cell line (in vitro resting and activated) and primary microglia models (primary resting and primary activated). CD9 = blue; CD81 = green; CD63 = red.

(**J**) Representative super resolution images of sEVs produced by BV2 cell line (in vitro). 100X objective with a 10X zoom factor. Scale bar = 200nm.

**Supplementary figure 3. TAMg-sEVs accelerate tumor growth and the infiltration of immune cells.**

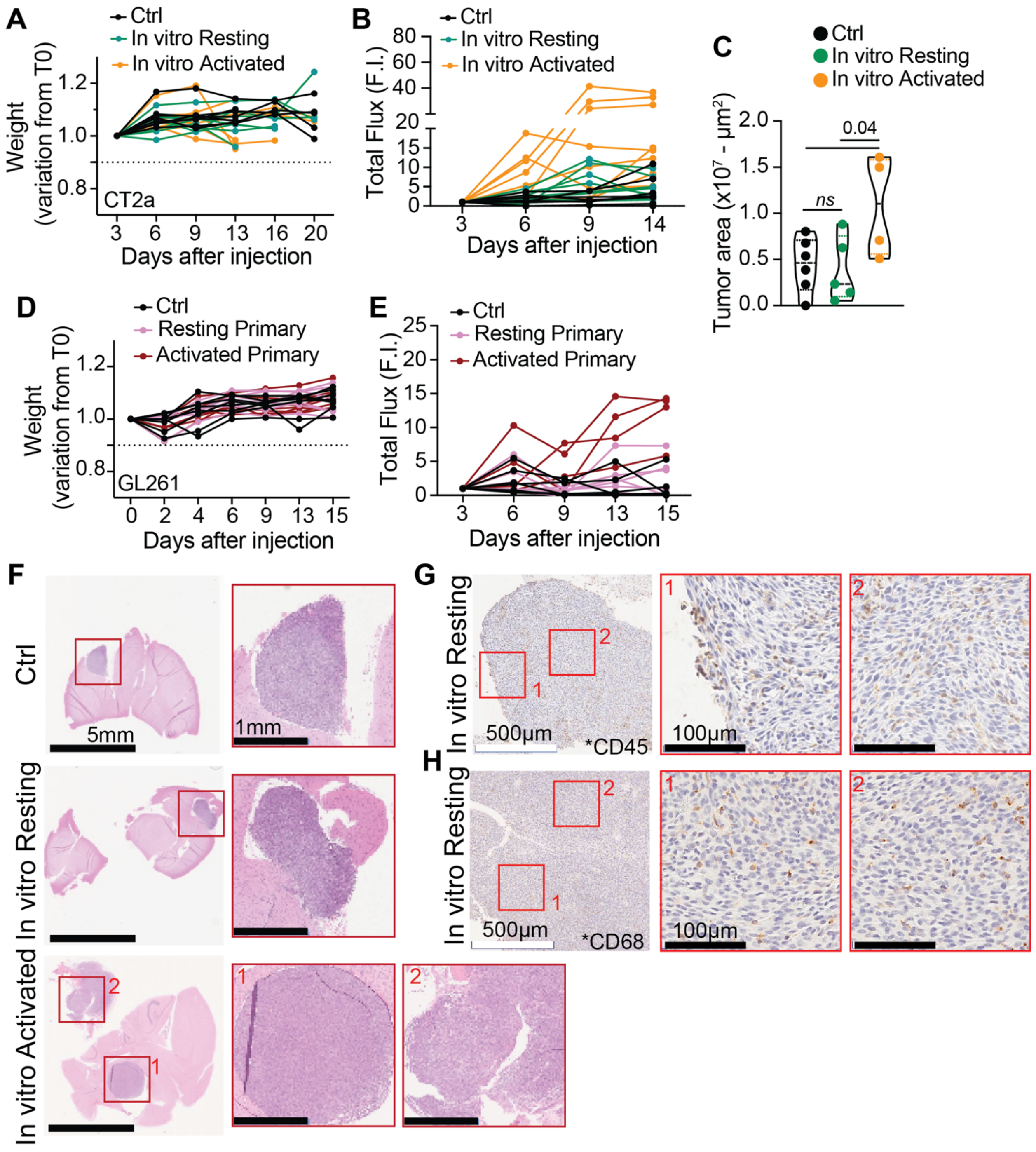


**(A-C)** CT2a glioma cell line was co-injected with BV2-sEVs (in vitro model) in C57BL/6 male and female albino mice. (A) Weight variation over time. (B) Tumor growth was tracked using an IVIS SpectrumCT In Vivo Imaging System at 3, 6, 9, and 14 days after injection. (C) Tumors were collected 14 days after the injection and tumor size confirmed by IHC. Ctrl = mice injected with

CT2a cells. In vitro Resting = mice co-injected with CT2a and resting BV2-derived sEVs. In vitro Activated = mice co-injected with CT2a and sEVs derived from BV2 cells activated with CT2a tumor supernatant. n= 8.

(**D-E**) GL261 glioma cell line was co-injected with primary microglia derived sEV in C57BL/6 female albino mice. Weight was monitored over time. (D) Weight variation over time. (E) Tumor growth was tracked using an IVIS SpectrumCT In Vivo Imaging System at 3, 6, 9, 13 and 15 days after injection. n= 5. Ctrl: mice injected with GL261. Primary Resting: mice co-injected with GL261 and sEVs isolated from resting primary microglia. Primary Activated: mice co-injected with GL261 and sEVs isolated from primary tumor associated microglia.

(**F-H**) The conditions are as in (A) and tumors were collected 14 days after the injection and stained for immunohistochemistry. (F) Representative H&E images of tumors from Ctrl and mice co-injected with sEVs from in vitro resting microglia and in vitro activated microglia. Red boxes = inset. (G-H) In vitro Resting microglia-sEVs condition were stained for CD45 (total immune cells, G) and CD68 (macrophages, H). Representative images, 4X objective, scale bar = 500µm. Red box = inset, 20X objective, scale bar = 100µm.

**Supplementary Figure 4. scRNA seq analysis of CD45+ cells infiltrating the tumor after TAMg-sEVs co-injection.**

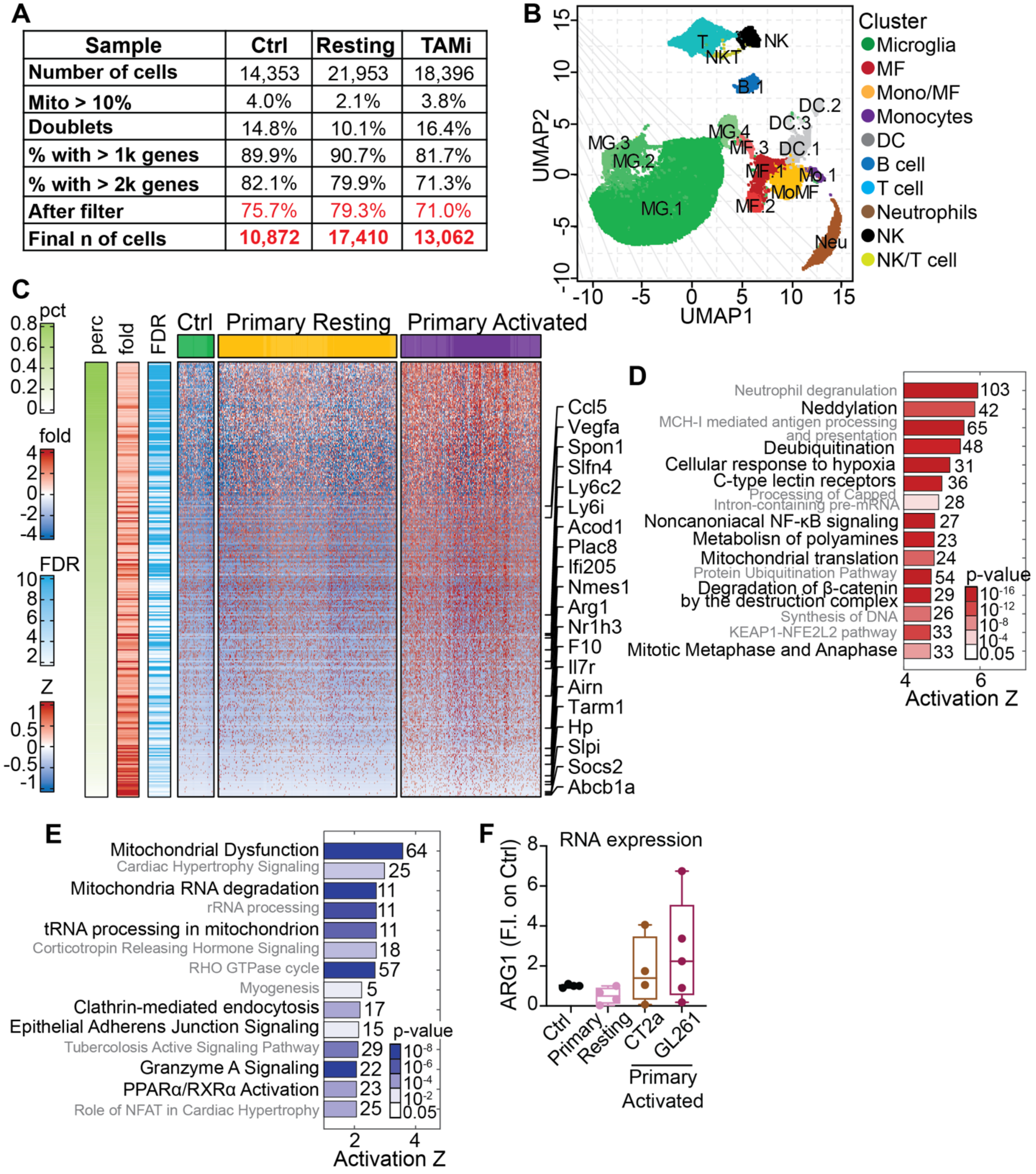


(A) scRNA seq data initial filtering for doublet, gene detection and mitochondrial content, resulted in at least 10k cells per sample.

(B) UMAP visualization of 18 clusters, annotated with assigned immune cell subtype information

(C) Expression heatmap of genes from most abundant macrophage cluster MF.1, significantly upregulated (FDR<5%) in mice co-injected with TAMi-sEVs (primary activated) compared to mice co-injected with resting microglia sEVs and Ctrl. Heatmap indicates expression Z-scores (Z). Genes are sorted by percent (pct, perc) of cells with detected expression. Top 20 most changed genes are highlighted

(D) Canonical pathways significantly activated in macrophages from the most abundant macrophage cluster MF.1 after TAMg-sEV injection, as identified by IPA enrichment analysis. In black are the pathways associated with macrophages activation within the tumor microenvironment.

(E) IPA canonical pathway enrichment analysis of TAMi-specific genes from the most abundant macrophage cluster MF.1. Top inhibited pathways in TAMi vs others.

(F) Macrophages were differentiated in the presence of sEVs for 8 days (as in figure 2D) and Arg1 expression was evaluated by qRT-PCR. Data are normalized to control (macrophages differentiated without sEVs.  Source of sEVs = primary resting and activated microglia. n=3. Source of macrophages = both male and female mice.

**Supplementary figure 5. Flow cytometry gating strategie.**

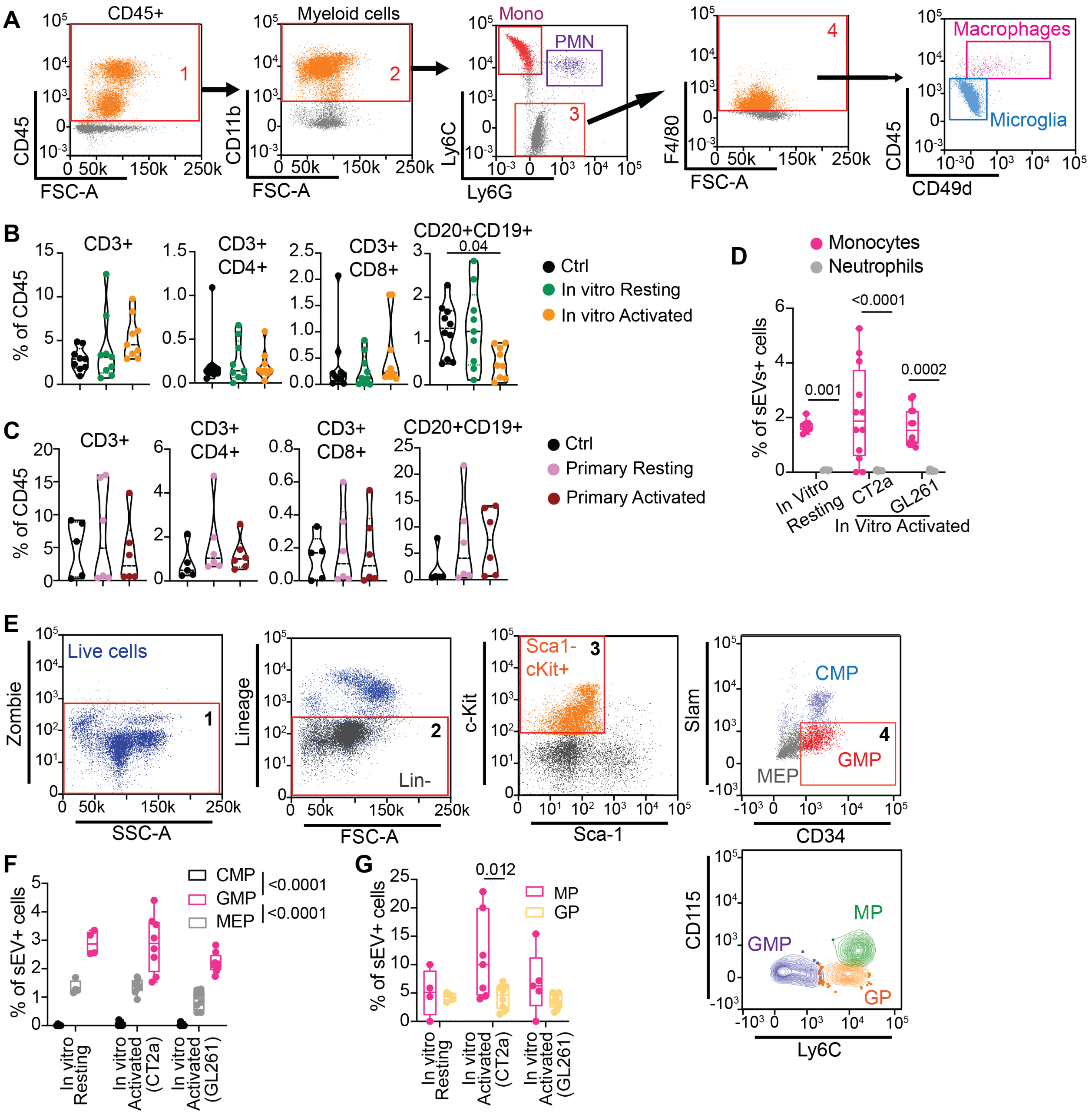


(**A**) Gating strategy for myeloid cells. Cells gated in single/live cells. 1 = CD45 positive population. 2 = CD11b positive cells in CD45. 3 = Ly6C and Ly6G neg in CD11b positive. Mono = monocytes, Ly6C positive and Ly6G negative cells in CD11b positive. PMN = neutrophils, Ly6G positive and Ly6C negative cells in CD11b positive. 4 = F4/80 positive cells in 3 gating. Macrophages = CD45 high CD49d high in gate 4. Microglia = CD45 low CD49d low in gate 4.

(**B-C**) CT2a or GL261 cells were co-injected with microglia-derived sEVs in C57BL/6 mice. (B) CT2a co-injected with in vitro sEVs. (C) GL261 co-injected with primary microglia-derived sEVs. Quantification of percentage in CD45+ cells of T cells (CD3+, CD4+ T cells, CD8+ T cells) and B cells (CD20+, CD19+). n=6. P-value = One-way ANOVA with Dunnett’s multiple comparison test.

(**D**) Cells were isolated from the BM of 8-week-old male and female naïve mice and co-cultured with DiD+ sEVs for 24 hours. sEVs internalization in myeloid cells was evaluated by flow cytometry. n= 3. p-value = Two-way ANOVA with Tukey's multiple comparisons test.

(**E**) Gating strategy for the identification of the different myeloid progenitor population in the BM. Live cells gated on single cells. CMP = Common Myeloid Progenitors. GMP = Granulocyte Myeloid Progenitors. MEP = Megakaryocyte-Erythroid Progenitors. MP = monocyte progenitors. GP = granulocyte progenitors.

(**F-G**) Cells were isolated from the BM of 8-week-old male and female naïve mice and co-cultured with DiD+ sEVs for 4 hours. (F) CMP = Common Myeloid Progenitors. GMP = Granulocyte Myeloid Progenitors. MEP = Megakaryocyte-Erythroid Progenitors. n=3. Šídák's multiple comparisons test. (G) MP = Monocyte Progenitors. GP = Granulocyte Progenitors. n= 3. p-value = Two-way ANOVA with Tukey's multiple comparisons test. Source of sEVs: BV2 cells.

**Supplementary Figure 6. TAMg-sEVs induce the expansion of highly motile inflammatory monocytes.**

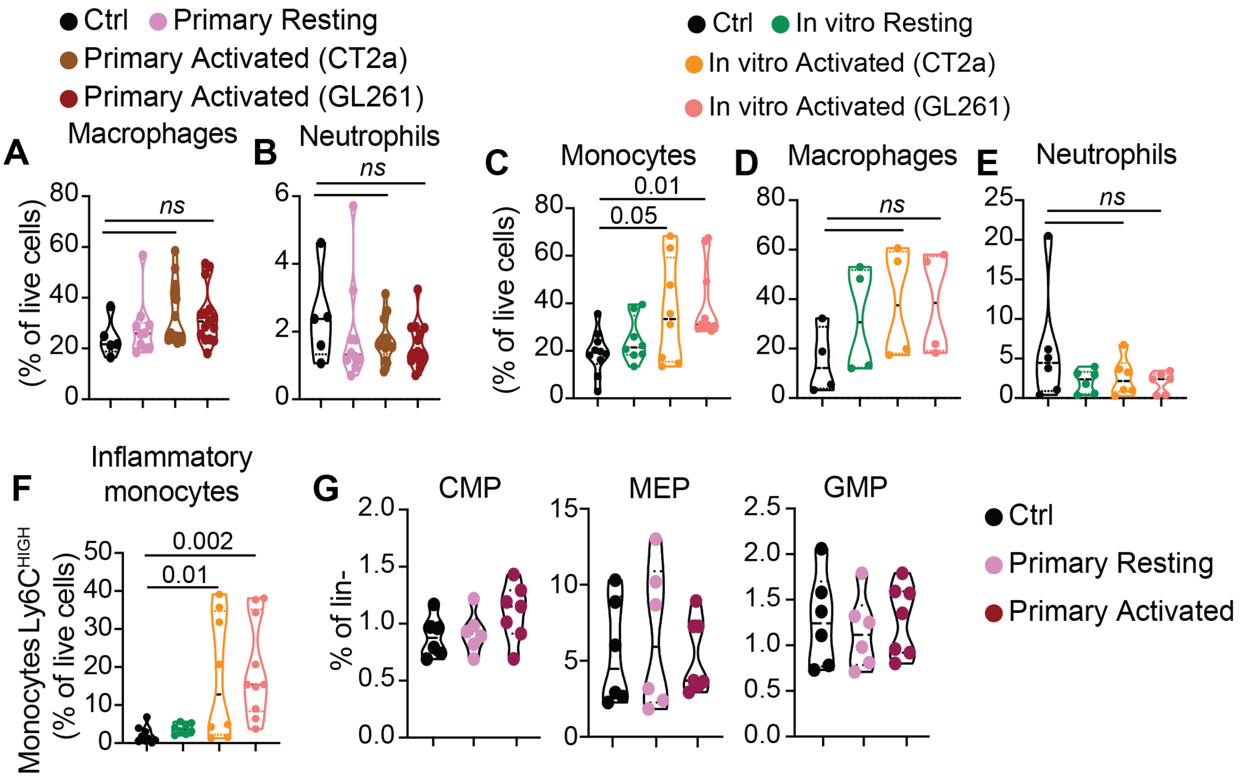


(**A-B**) Immune cells isolated from the bone marrow of C57BL/6 male and female naïve mice were co-cultured for 6 days with sEVs produced by primary resting microglia or primary tumor-associated microglia (isolated from CT2a or GL261 injected mice). The percentage of macrophages (A) and neutrophils (B) was measured at day 6 by flow cytometry. n = 6. p-value = One-way ANOVA with Dunnett's multiple comparisons test.

(**C-E**) The conditions are as in (A) and immune cells were co-cultured with sEVs produced by BV2 cells. In vitro Resting: BV2 not activated. CT2a: BV2 activated with CT2a tumor supernatant. GL261: BV2 activated with GL261 tumor supernatant. The percentage of monocytes (C), macrophages (D) and neutrophils (E) was measured by flow cytometry. n = 5. p-value = One-way ANOVA with Dunnett's multiple comparisons test.

(**F**) The conditions are as in (C) and the percentage of inflammatory monocytes ($Ly6C^{HIGH}$) was measured by flow cytometry. n = 5. p-value = One-way ANOVA with Dunnett's multiple comparisons test.

(**G**) Microglia-derived sEVs were co-injected with GL261 in 8-week-old naïve female mice and the percentage of Common Myeloid Progenitors (CMP), Megakaryocyte-Erythroid Progenitor (MEP) and Granulocyte-Monocyte Progenitor (GMP) was evaluated by flow cytometry after 3 days after the injection in both spine and femur BM. n=6.

**Supplementary Figure 7. TAMg-sEVs activate CCR2$^{HIGH}$ motile monocytes in the bone marrow.**

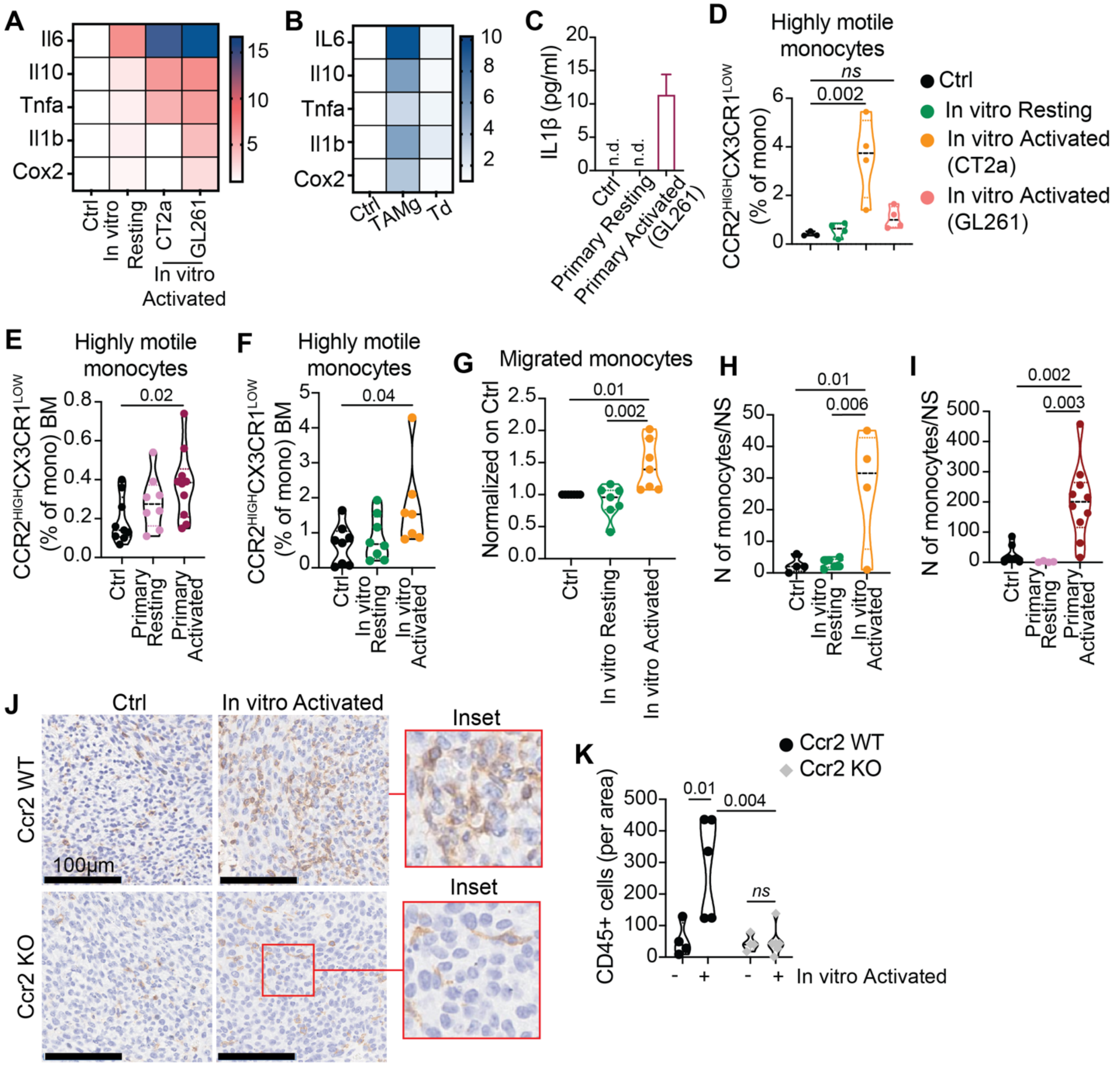


(**A**) Immune cells were isolated from the bone marrow of male and female tumor-free mice and co-cultured with BV2-derived sEVs for six days. Changes in gene expression was evaluated by qRT-PCR. n= 6.

(**B**) The conditions are as in (A) and immune cells were co-cultured with primary microglia-derived (TAMg) and tumor-derived (Td) sEVs for 6 days. Differences in gene expression were evaluated by qRT-PCR. n=6.

(**C**) The conditions are as in (A) and immune cells were co-cultured with sEVs produced by primary microglia. The production of the cytokine IL-1β was tested at day 6 by ELISA. n.d. = not detected. n=3.

(**D**) The conditions are as in (A) and the percentage highly motile inflammatory monocytes ($CCR2^{HIGH}CX3CR1^{LOW}$) was measured by flow cytometry at day 6. n = 5. p-value = One-way ANOVA with Dunnett's multiple comparisons test.

(**E-F**) Microglia-derived sEVs were co-injected with GL261 (E) or CT2a (F) in 8-week-old naïve female mice and the percentage of highly motile monocytes in the femur bone marrow was evaluated by flow cytometry 7 days after the injection. n=6. p-value = One-way ANOVA with Dunnett's multiple comparisons test.

(**G**) Immune cells were isolated from the bone marrow of male and female tumor-free mice and co-cultured with BV2-derived sEVs for six days. At day 6 monocytes were collected, counted and seeded on the upper part of an 8μm transwell insert. The number of migrated cells (bottom) was counted and normalized on control conditions. n=3. p-value = 1way ANOVA with Dunnett's multiple comparison test.

(**H-I**) Neurospheres (NS) generated from tumor bearing mice were co-cultured for 24 hours with monocytes conditioned with microglia-derived sEVs (as in G). The number of monocytes infiltrating the NS was quantified. (H) Myeloid cells co-cultured with sEVs isolated from BV2 cells. (**I**) Myeloid cells co-cultured with primary microglia-derived sEVs. p-value = 1way Anova with Dunnett's multiple comparisons test.

(**J-L**) C57BL/6 WT (Ccr2 WT) and Ccr2-deficient (Ccr2 KO) female mice were co-injected with CT2a cell line and BV2 derived sEVs. Brains were collected at day 13 and processed by immunohistochemistry. (**J**) Representative images of CD45 staining. Objective = 20X. (**K**) Quantification of CD45+ cells per 10X area. n= 4. Two-way ANOVA with Turkey's multiple comparison test.

### SUPPLEMENTARY TABLES

**Table S1. BV2 activation: mRNA expression normalized on resting microglia**

| | Primers | | |
|---|---|---|---|
| **Samples** | **Tgf-β** | **Arg1** | **Il10** |
| **In vitro Activated GL261-1** | 2.409 | 0.3 | 10.8 |
| **In vitro Activated GL261-2** | 3.08 | 5.09 | 9.7 |
| **In vitro Activated GL261-3** | 1.232 | 2.34 | 11.9 |
| **In vitro Activated GL261-4** | 1.346 | na | 1.8 |
| **In vitro Activated GL261-5** | 1.862 | na | 1.3 |
| **In vitro Activated GL261-6** | 1.398 | na | 1.95 |
| **In vitro Activated GL261-7** | 1.358 | na | 1.5 |
| **In vitro Activated CT2a-1** | 1.5 | 383.5 | 0.6 |
| **In vitro Activated CT2a-2** | 1.1 | 1.9 | 1.3 |
| **In vitro Activated CT2a-3** | 3.6 | 1.7 | 2.8 |
| **In vitro Activated CT2a-4** | 1.3 | 2.4 | 1.4 |
| **In vitro Activated CT2a-5** | 1.5 | na | 1 |
| **In vitro Activated CT2a-6** | 1 | 1.5 | 1.5 |
| **In vitro Activated CT2a-7** | 4.8 | 122.8 | 30.5 |
| **In vitro Activated CT2a-8** | 4.8 | 110.1 | 30.5 |

**Table S2. scRNAseq_ClusterClassification**

| cluster | cells n | cells % | cell type Primary | secondary (% cluster) |
|---|---|---|---|---|
| **MG.1** | 22,806 | 59.50% | Microglia | |
| **MG.2** | 2,052 | 5.40% | Microglia | MF (11%) |
| **MG.3** | 862 | 2.20% | Microglia | |
| **MG.4** | 823 | 2.10% | Microglia | MF (9%) |
| **MF.1** | 1,347 | 3.50% | MF | |
| **MF.2** | 657 | 1.70% | MF | |
| **MF.3** | 295 | 0.80% | MF | MG (11%) |
| **MoMF** | 1,432 | 3.70% | Monocyte | MF (31%) |
| **Mo** | 338 | 0.90% | Monocyte | |
| **DC.1** | 924 | 2.40% | DC | Mono (11%), MF (10%) |
| **DC.2** | 295 | 0.80% | DC | |
| **DC.3** | 288 | 0.80% | DC | |
| **B** | 877 | 2.30% | B cell | |
| **Neu** | 1,746 | 4.60% | Neutrophil | |
| **NK** | 776 | 2.00% | NK cell | |
| **T** | 2,637 | 6.90% | T cell | NK (7%) |
| **NKT** | 167 | 0.40% | T cell | NK (35%) |

**Table S3. scRNAseq_Cluster/Cell enrichment: Mice co-injected with Primary Activated microglia-derived sEVs vs Ctrl and primary resting microglia-sEVs**

| group | name | GAMi n cells | GAMi ratio vs c+R | ctrl | ReMi | GAMi | ctrl | ReMi | GAMi |
|---|---|---|---|---|---|---|---|---|---|
| C\|T | MG.2\|Macrophages | 213 | 14.5 | | | | 2 | 20 | 213 |
| C\|T | NKT\|T cells | 75 | 4.1 | | | | 9 | 25 | 75 |
| clust | NKT | 111 | 3.9 | | | | 17 | 39 | 111 |
| clust | DC.3 | 189 | 3.6 | | | | 27 | 72 | 189 |
| C\|T | DC.3\|DC | 189 | 3.6 | | | | 27 | 72 | 189 |
| C\|T | NKT\|NK cells | 36 | 3.5 | | | | 8 | 14 | 36 |
| C\|T | MoMF\|Monocytes | 621 | 3.1 | | | | 99 | 272 | 621 |
| C\|T | DC.1\|DC | 415 | 2.8 | | | | 109 | 203 | 415 |
| type | DC | 754 | 2.6 | | | | 162 | 394 | 754 |
| type | Monocytes | 810 | 2.6 | | | | 195 | 431 | 810 |
| clust | DC.1 | 496 | 2.3 | | | | 134 | 294 | 496 |
| C\|T | DC.1\|Monocytes | 61 | 2.2 | | | | 7 | 38 | 61 |
| clust | MoMF | 771 | 2.1 | | | | 152 | 509 | 771 |
| clust | DC.2 | 150 | 1.7 | | | | 26 | 119 | 150 |
| C\|T | DC.2\|DC | 150 | 1.7 | | | | 26 | 119 | 150 |
| clust | MG.2 | 897 | 1.4 | | | | 518 | 637 | 897 |
| C\|T | MF.3\|Macrophages | 102 | 1.3 | | | | 64 | 98 | 102 |
| clust | Neu | 754 | 1.3 | | | | 187 | 805 | 754 |
| type | Neutrophils | 754 | 1.3 | | | | 187 | 805 | 754 |
| C\|T | Neu\|Neutrophils | 754 | 1.3 | | | | 187 | 805 | 754 |
| type | T cells | 1105 | 1.3 | | | | 262 | 1190 | 1105 |
| type | Macrophages | 1270 | 1.3 | | | | 459 | 1378 | 1270 |
| type | NK cells | 417 | 1.2 | | | | 146 | 460 | 417 |
| C\|T | T\|T cells | 1030 | 1.2 | | | | 253 | 1165 | 1030 |
| clust | NK | 317 | 1.2 | | | | 99 | 360 | 317 |
| C\|T | NK\|NK cells | 317 | 1.2 | | | | 99 | 360 | 317 |
| clust | T | 1094 | 1.2 | | | | 292 | 1251 | 1094 |
| clust | Mo.1 | 128 | 1.2 | | | | 89 | 121 | 128 |
| C\|T | Mo.1\|Monocytes | 128 | 1.2 | | | | 89 | 121 | 128 |
| C\|T | MG.4\|Macrophages | 32 | 1.2 | | | | 4 | 37 | 32 |
| clust | MF.3 | 108 | 1.1 | | | | 78 | 109 | 108 |
| C\|T | MG.2\|Microglia | 684 | 1.1 | | | | 516 | 617 | 684 |
| clust | MF.1 | 532 | 1.1 | | | | 137 | 678 | 532 |
| C\|T | MF.1\|Macrophages | 532 | 1.1 | | | | 137 | 678 | 532 |
| C\|T | T\|NK cells | 64 | 1 | | | | 39 | 86 | 64 |
| clust | MF.2 | 221 | 1 | | | | 181 | 255 | 221 |
| C\|T | MF.2\|Macrophages | 221 | 1 | | | | 181 | 255 | 221 |
| clust | MG.3 | 360 | 0.9 | | | | 313 | 189 | 360 |
| C\|T | MG.3\|Microglia | 360 | 0.9 | | | | 313 | 189 | 360 |
| clust | MG.4 | 252 | 0.9 | | | | 193 | 378 | 252 |
| C\|T | MG.4\|Microglia | 220 | 0.9 | | | | 189 | 341 | 220 |
| C\|T | MoMF\|Macrophages | 150 | 0.9 | | | | 53 | 237 | 150 |

| | | | | | | | | | |
|---|---|---|---|---|---|---|---|---|---|
| type | Microglia | 6683 | 0.7 | | | | 8367 | 11216 | 6683 |
| clust | MG.1 | 5413 | 0.6 | | | | 7335 | 10058 | 5413 |
| C\|T | MG.1\|Microglia | 5413 | 0.6 | | | | 7335 | 10058 | 5413 |
| clust | B.1 | 235 | 0.6 | | | | 105 | 537 | 235 |
| type | B cells | 235 | 0.6 | | | | 105 | 537 | 235 |
| C\|T | B.1\|B cells | 235 | 0.6 | | | | 105 | 537 | 235 |
| C\|T | DC.1\|Macrophages | 20 | 0.5 | | | | 18 | 53 | 20 |
| C\|T | MF.3\|Microglia | 6 | 0.4 | | | | 14 | 11 | 6 |

**Table S4. Differentially expressed genes in different cluster. Mice co-injected with Primary Activated microglia-derived sEVs vs Ctrl and primary resting microglia-sEVs.**

| focus set | cell group | pvalue<0.05 | | | FDR<5% | | |
|---|---|---|---|---|---|---|---|
| | | all | up | down | all | up | dn |
| | MG.2\|Microglia | **5198** | 860 | 4338 | **1994** | 612 | 1382 |
| **MG** | MG.1\|Microglia | **1748** | 1085 | 663 | **910** | 564 | 346 |
| **MFX** | MG.2\|Macrophages | **1464** | 353 | 1111 | **108** | 82 | 26 |
| **MF** | MF.1\|Macrophages | **1431** | 1002 | 429 | **534** | 369 | 165 |
| | Neu\|Neutrophils | **1241** | 417 | 824 | **590** | 264 | 326 |
| | T.1\|T cells | **801** | 548 | 253 | **350** | 237 | 113 |
| **MoX** | MoMF\|Monocytes | **726** | 118 | 608 | **261** | 64 | 197 |
| | MG.3\|Microglia | **667** | 195 | 472 | **276** | 137 | 139 |
| | MG.4\|Microglia | **590** | 230 | 360 | **142** | 76 | 66 |
| | MoMF\|Macrophages | **581** | 320 | 261 | **156** | 68 | 88 |
| | DC.1\|DC | **550** | 290 | 260 | **197** | 88 | 109 |
| | MF.3\|Macrophages | **521** | 106 | 415 | **165** | 46 | 119 |
| | NK\|NK cells | **349** | 253 | 96 | **89** | 74 | 15 |
| | B.1\|B cells | **344** | 229 | 115 | **107** | 84 | 23 |
| **Mo** | Mo.1\|Monocytes | **239** | 137 | 102 | **26** | 22 | 4 |
| | DC.1\|Monocytes | **237** | 111 | 126 | **27** | 25 | 2 |
| | DC.1\|Macrophages | **235** | 136 | 99 | **1** | 1 | 0 |
| | MF.2\|Macrophages | **181** | 77 | 104 | **9** | 8 | 1 |
| | DC.3\|DC | **170** | 42 | 128 | **18** | 1 | 17 |
| | NKT\|T cells | **165** | 43 | 122 | **46** | 16 | 30 |
| | DC.2\|DC | **144** | 86 | 58 | **19** | 11 | 8 |
| | NKT\|NK cells | **132** | 39 | 93 | **10** | 9 | 1 |
| | T.1\|NK cells | **121** | 82 | 39 | **0** | 0 | 0 |
| | MF.3\|Microglia | **46** | 34 | 12 | **2** | 2 | 0 |
| | MG.4\|Macrophages | **33** | 5 | 28 | **1** | 1 | 0 |

**Table S5. Flow cytometry panels.**

| Ex-vivo | | | |
|---|---|---|---|
| ***Microglia*** | | ***Myeloid progenitors*** | |
| **Antibody** | ***fluorochrome*** | **Antibody** | ***fluorochrome*** |
| FC block (CD16/CD32) | unconj | FC block (CD16/CD32) | unconj |
| UV Zombie | UV447 | UV Zombie | UV447 |
| CD11b | BV785 | CD115 | BV421 |
| F4/80 | PercP-Cy5.5 | Lineage | Pacific Bue |
| CD206 | PE-Fire 700 | Ly6C | BV786 |
| TMEM119 | PeCy7 | CD34 | FITC |
| ARG1 | APC | Ly-6A/E (Sca-1) | PE |
| ***Myeloid cells*** | | CD150 (SLAM) | PE/Cy7 |
| **Antibody** | ***fluorochrome*** | **sEV** | **DiD (APC)** |
| FC block (CD16/CD32) | unconj | CD117 (c-kit) | APC/Cy7 |
| UV Zombie | UV447 | | |
| CD115 | BV421 | | |
| CD163 | BV605 | | |
| CD11C | BV711 | | |
| CD11b | BV785 | | |
| LY6G | FITC | | |
| ClassII | PerCP/Cyanine5.5 | | |
| F480 | PE | | |
| CX3CR1 | PE-Dazzle-594 | | |
| CCR2 | PE-Fire 640 | | |
| Ly6C | PE-Cy7 | | |
| **sEV** | **DiD (APC)** | | |
| *red = intracellular | | | |

| In vivo | | | |
|---|---|---|---|
| **BRAIN** | | **Bone Marrow** | |
| ***Micro/Macro*** | | ***Myeloid cells*** | |
| **Antibody** | ***fluorochrome*** | **Antibody** | ***fluorochrome*** |
| FC block (CD16/CD32) | unconj | FC block (CD16/CD32) | unconj |
| UV Zombie | UV447 | UV Zombie | UV447 |
| ARG1 | BV421 | CD115 | BV421 |
| Ly-6C/Ly6G | Pacific Blue | CD163 | BV605 |
| CD45 | BV650 | CD11C | BV711 |
| CD11c | BV711 | CD11b | BV785 |
| CD11b | BV786 | LY6G | FITC |
| CD49d | AF488 | ClassII | PerCP/Cyanine5.5 |
| F4/80 | PerCP-Cy5.5 | F480 | PE |
| CX3CR1 | PE-DAZZLE-594 | CX3CR1 | PE-Dazzle-594 |
| iNOS | PE-Cy5 | CCR2 | PE-Fire 640 |
| CD206 | PE-Fire 700 | Ly6C | PE-Cy7 |
| sEV | DiD (APC) | **sEV** | **DiD (APC)** |
| CD86 | AF700 | ***Myeloid progenitors*** | |
| CD163 | APC-Cy7 | **Antibody** | ***fluorochrome*** |

| DCs-Myeloid mix | |
|---|---|
| **Antibody** | ***fluorochrome*** |
| FC block (CD16/CD32) | unconj |
| UV Zombie | UV447 |
| CD11B | BV421 |
| CD45 | BV650 |
| CD11c | BV710 |
| Ly6C | BV786 |
| Ly6g | FITC |
| Class II | PerCP-Cy5.5 |
| F4/80 | PE |
| **sEV** | **DiD (APC)** |

| T cells mix | |
|---|---|
| **Antibody** | ***fluorochrome*** |
| FC block (CD16/CD32) | unconj |
| UV Zombie | UV447 |
| CD8 | BV421 |
| CD45 | BV650 |
| CD4 | BV711 |
| CD19 | BV750 |
| CD69 | BV786 |
| CD3 | PE |
| CD20 | PE-Cy7 |

| | |
|---|---|
| FC block (CD16/CD32) | unconj |
| UV Zombie | UV447 |
| CD115 | BV421 |
| Lineage | Pacific Bue |
| Ly6C | BV786 |
| CD34 | FITC |
| Ly-6A/E (Sca-1) | PE |
| CD150 (SLAM) | PE/Cy7 |
| **sEV** | **DiD (APC)** |
| CD117 (c-kit) | APC/Cy7 |

| T cells mix | |
|---|---|
| **Antibody** | ***fluorochrome*** |
| FC block (CD16/CD32) | unconj |
| UV Zombie | UV447 |
| CD8 | BV421 |
| CD45 | BV650 |
| CD4 | BV711 |
| CD19 | BV750 |
| CD69 | BV786 |
| CD3 | PE |
| CD20 | PE-Cy7 |

*red = intracellular

**Table S6. Flow cytometry, bone marrow. 7 days post injection**

| | Injection CT2a | | | Injection GL261 | | |
|---|---|---|---|---|---|---|
| | Ctrl | In vitro Resting | In vitro Activated | Ctrl | Primary Resting | Primary Activated |
| CD3+ | 3.59 | 6.04 | 5.32 | 1.68 | 0.81 | 1.06 |
| | 4.46 | 5.42 | 6.68 | 1.47 | 1.16 | 0.45 |
| | 5.11 | 4.55 | 5.91 | 1.32 | 0.89 | |
| | 10.3 | 9.47 | 14.6 | 2.53 | 1.89 | 3.03 |
| | 11.5 | 8.01 | 11.8 | 1.6 | 1.88 | 1.96 |
| | 1.78 | 1.6 | 13.4 | 1.07 | 3.59 | 1.86 |
| | 3.2 | 1.94 | 1.88 | 1.83 | 0.78 | 1.25 |
| | 3.15 | 3.14 | 3.09 | 1.55 | 1.27 | 0.28 |
| | | | 2.85 | 1.36 | 1.13 | |
| CD3+ CD4+ | 0.45 | 0.97 | 0.86 | 1.42 | 0.69 | 0.98 |
| | 0.47 | 0.81 | 1.23 | 1.23 | 1.02 | 0.38 |
| | 0.75 | 0.6 | 0.89 | 1.07 | 0.81 | |
| | 0.38 | 0.36 | 0.32 | 0.77 | 0.6 | 0.87 |
| | 0.46 | 0.42 | 0.56 | 0.42 | 0.5 | 0.61 |
| | 0.066 | 0.25 | 0.3 | 0.32 | 1 | 0.5 |
| | 0.046 | 0.078 | 0.067 | 0.43 | 0.19 | 0.31 |
| | 0.13 | 0.099 | 0.079 | 0.32 | 0.52 | 0.087 |
| | | 0.11 | 0.12 | 0.34 | 0.34 | |
| CD3+ CD8+ | 0.73 | 1.94 | 1.81 | 0.48 | 0.16 | 0.4 |
| | 0.71 | 2.03 | 2.56 | 0.42 | 0.22 | 0.043 |
| | 1.53 | 1.43 | 2.01 | 0.36 | 0.27 | |
| | 0.24 | 0.21 | 0.17 | 1.29 | 0.86 | 1.46 |
| | 0.37 | 0.42 | 0.48 | 0.8 | 0.94 | 0.97 |
| | 0.026 | 0.17 | 0.19 | 0.37 | 1.91 | 0.87 |
| | 0.022 | 0.05 | 0.034 | 0.51 | 0.18 | 0.5 |
| | 0.05 | 0.062 | 0.061 | 0.42 | 0.23 | 0.041 |
| | | 0.048 | 0.049 | 0.31 | 0.36 | |
| CD20+ CD19+ | 18.5 | 28.5 | 27.1 | 5.8 | 2.08 | 1.98 |
| | 25.1 | 32 | 34.3 | 5.02 | 2.61 | 1.61 |
| | 21.8 | 29.7 | 30.7 | 4 | 2.28 | |
| | 10.4 | 14.3 | 8.41 | 23.1 | 28.7 | 21.6 |
| | 12.7 | 13.8 | 12.7 | 25.6 | 29.3 | 27.9 |
| | 17.9 | 13.3 | 13.7 | 21.9 | 22.1 | 16.4 |
| | 14.9 | 15.9 | 16.6 | 16.3 | 14.4 | 19.9 |
| | 7.09 | 16.9 | 13.8 | 10.5 | 19 | 4.28 |
| | | 15.4 | 16.1 | 12.8 | 11.7 | |
| PMN | 26.2 | 18.8 | 17.8 | 29.1 | 29.9 | 26.6 |
| | 24.4 | 19.3 | 15.7 | 31.2 | 27.4 | 31.2 |
| | 24.6 | 25.5 | 15.5 | 30.7 | 27.8 | 27 |
| | 2.86 | 2.47 | 4.65 | 32.9 | 39.1 | 42.1 |
| | 0.5 | 1.36 | 2.05 | 32.9 | 39.4 | 31.8 |
| | 2.98 | 1.9 | 2.2 | 28.5 | 32.6 | 23.1 |

| | | | | | | |
|---|---|---|---|---|---|---|
| | | | | 38.1 | 44.6 | 47.9 |
| | | | | 38.7 | 44.3 | 29.6 |
| | | | | 33.5 | 38.2 | |
| | 0.1 | 0.13 | 0.12 | 0.038 | 0.025 | 0.067 |
| | 0.15 | 0.095 | 0.12 | 0.037 | 0.04 | 0.032 |
| | 0.11 | 0.1 | 0.16 | 0.032 | 0.036 | 0.048 |
| | 0.35 | 0.33 | 0.35 | 0.7 | 0.66 | 0.59 |
| DCs | 0.29 | 0.42 | 0.25 | 0.55 | 0.63 | 0.69 |
| | 0.4 | 0.3 | 0.29 | 0.64 | 0.4 | 0.89 |
| | | | | 0.17 | 0.25 | 0.22 |
| | | | | 0.24 | 0.23 | 0.25 |
| | | | | 0.19 | 0.23 | |

**Table S7. List of flow cytometry antibodies.**

| **Suppl.Table7_List of flow cytometry antibodies** | | | | |
|---|---|---|---|---|
| **Target protein** | **Fluorochrome** | **Company** | **Catalog number** | **Clone** |
| ARG1 | APC | Invitrogen | 17-3697-82 | A1exF5 |
| ARG1 | BV421 | Invitrogen | 404-3697-80 | A1exF5 |
| CCR2 | PE-Fire 640 | BioLegend | 150639 | SA203G11 |
| CD115 | BV421 | BioLegend | 135513 | AFS98 |
| CD117 (c-kit) | APC/Cy7 | BioLegend | 105825 | 2B8 |
| CD11b | BV785 | BioLegend | 101243 | M1/70 |
| CD11B | BV421 | BioLegend | 101236 | M1/70 |
| CD11C | BV711 | BioLegend | 117349 | N418 |
| CD150 (SLAM) | PE/Cy7 | BioLegend | 115913 | TC15-12F12.2 |
| CD163 | BV605 | BioLegend | 155327 | S15049I |
| CD163 | APC-Cy7 | BioLegend | 155324 | S15049I |
| CD19 | BV750 | BD Bioscience | 747332 | 1D3 |
| CD20 | PE-Cy7 | Invitrogen | 25-0452-81 | RA3-6B2 |
| CD206 | PE-Fire 700 | BioLegend | 141741 | C068C2 |
| CD206 | APC-eFluor 780 | Invitrogen | 47-2061-82 | MR6F3 |
| CD3e | PE | BD Bioscience | 553063 | 145-2C11 |
| CD34 | FITC | Invitrogen | 11-0341-82 | RAM34 |
| CD4 | BV711 | BioLegend | 100447 | GK1.5 |
| CD45 | BV650 | BioLegend | 103151 | 30-F11 |
| CD49d | AF488 | BioLegend | 103611 | R1-2 |
| CD64 | FITC | BioLegend | 139316 | X54-5/7.1 |
| CD69 | BV785 | BioLegend | 104543 | H1.2F3 |
| CD8a | BV421 | BioLegend | 100753 | 53-6.7 |
| CD86 | AF700 | BioLegend | 105024 | GL-1 |
| ClassII | PerCP/Cyanine5.5 | BioLegend | 116415 | AF6-120.1 |
| CX3CR1 | PE-Dazzle-594 | BioLegend | 149013 | SA011F11 |
| F4/80 | PercP-Cy5.5 | BD Bioscience | 567202 | T45-2342 |
| F4/80 | PE | BioLegend | 123110 | BM8 |
| iNOS | PE-Cy.5 | Invitrogen | 35-5920-80 | CXNFT |
| Lineage | Pacific Bue | BioLegend | 133305 | 17A2; RB6-8C5; RA3-6B2; Ter-119; M1/70 |
| Ly-6A/E (Sca-1) | PE | BioLegend | 160905 | W18174A |
| Ly-6C/Ly6G | Pacific Blue | BioLegend | 108429 | RB6-8C5 |
| Ly-6C | PE-Cy7 | BD Bioscience | 560593 | AL-21 |
| Ly-6C | BV786 | BD Bioscience | 569012 | AL-21 |
| Ly-6G | FITC | BD Bioscience | 551460 | 1A8 |

| | | | | |
|---|---|---|---|---|
| TMEM119 | PeCy7 | Invitrogen | 25-6119-80 | V3RT1GOsz |
| TNFα | BV650 | BD Bioscience | 563943 | MP6-XT22 |

**Table S8. List of qRT-PCR primers.**

| Gene name | Sequence (5’ – 3’) |
|---|---|
| m_18s_FW | CGGAAAATAGCCTTCGCCATCAC |
| m_18s_RV | ATCACTCGCTCCACCTCATCCT |
| m-TGF-β_FW | TGATACGCCTGAGTGGCTGTCT |
| m-TGF-β_RV | CACAAGAGCAGTGAGCGCTGAA |
| m-iNOS_FW | GAGACAGGGAAGTCTGAAGCAC |
| m-iNOS_RV | CCAGCAGTAGTTGCTCCTCTTC |
| m_COX2_FW | GCGACATACTCAAGCAGGAGCA |
| m_COX2_RV | AGTGGTAACCGCTCAGGTGTTG |
| m_IL-10_RV | CGGTTAGCAGTATGTTGTCCAGC |
| m_IL-10_FW | CGGGAAGACAATAACTGCACCC |
| m_TNFα_RV | GCCATAGAACTGATGAGAGGGAG |
| m_TNFα_FW | GGTGCCTATGTCTCAGCCTCTT |
| m_IL-6_RV | CTGCAAGTGCATCATCGTTGTTC |
| m_IL-6_FW | TACCACTTCACAAGTCGGAGGC |
| m_IL-1β_RV | CGGTTAGCAGTATGTTGTCCAGC |
| m_IL-1β_FW | CGGGAAGACAATAACTGCACCC |
| m_Arginase1_RV | GAATCCTGGTACATCTGGGAAC |
| m_Arginase1_FW | GAATCTGCATGGGCAACC |
